# Numerous independent gains of torpor and hibernation across endotherms, linked with adaptation to diverse environments

**DOI:** 10.1101/2023.12.12.571278

**Authors:** Dimitrios - Georgios Kontopoulos, Danielle L. Levesque, Michael Hiller

## Abstract

Many endotherms from diverse taxonomic groups can respond to environmental changes through dormancy, i.e., by greatly reducing their energy expenditure for up to 24 hours (daily torpor) or longer (hibernation). We currently have a poor understanding of how dormancy evolved across endotherms and its associations with physiological traits and ecological factors. To fill this gap, we thoroughly examine the evolutionary patterns of dormancy and its links with 21 key ecophysiological variables across 1,338 extant endotherms. We find that daily torpor and hibernation are parts of a dormancy continuum, and that there are several, albeit weak, associations between dormancy and species’ physiological or environmental characteristics. Furthermore, we show that early endotherm ancestors likely did not hibernate and that this trait evolved multiple times in independent lineages. Overall, our results provide an explanation for the remarkable variation in dormancy patterns, even among species occupying highly similar niches.

## Introduction

Whole-body endotherms (mammals and birds) are able to maintain their body temperature at a high and relatively constant value across a wide range of ambient temperatures by means of a much higher resting metabolic rate than that of ectotherms (Grigg *et al*. 2022). The energy expenditure of a resting endotherm is at its lowest and constant over a range of temperatures called the thermoneutral zone, but many endotherm species live below that zone and, therefore, spend significant amounts of energy thermoregulating (Clarke 2017; Geiser 2021). Elevated energy costs can be unsustainable under challenging environmental conditions such as extreme temperatures, or food or water shortages. In response to such challenges, numerous mammals and birds undergo dormancy, which involves programmed and reversible decreases in body temperature and metabolic rate (Geiser 2021; Nowack *et al*. 2020; Ruf & Geiser 2015). Dormancy patterns of endotherms can be broadly classified based on their duration as either a) daily torpor or b) prolonged torpor or hibernation (but see Levesque *et al*. 2023). The former lasts for less than 24 hours, whereas the latter can last from a day up to more than a year, albeit typically interrupted by brief rewarming periods (Geiser 2021; Nowack *et al*. 2020). Differentiating between prolonged torpor (multi-day torpor bouts sensu Dausmann 2014) and hibernation is less straightforward (e.g., see Nowack *et al*. 2020) and we henceforth refer to dormancy patterns that last for more than 24 hours as hibernation for the sake of brevity. Another key difference between daily torpor and hibernation is that the former results in less pronounced decreases in body temperature and metabolic rate than the latter (Geiser 2021). It is worth noting that some species also use dormancy for other purposes besides energy conservation, such as for predator avoidance or for coexistence with competitors (e.g., see Bieber & Ruf 2009; Levy *et al*. 2011; Powers 2004).

In mammals, daily torpor and hibernation have been observed in most large orders, including in placentals, marsupials, and monotremes (Geiser 2021; Lovegrove 2012; Nowack *et al*. 2020; Ruf & Geiser 2015). In contrast, many orders of birds include species that are capable of daily torpor (e.g., Apodiformes, Passeriformes, Columbiformes, Strigiformes), but only a single bird species—the common poorwill (*Phalaenoptilus nuttallii*)—is known to hibernate (Geiser 2021; McKechnie *et al*. 2023; McNab 2009; Ruf & Geiser 2015; Woods *et al*. 2019). Hence, dormancy as a key survival strategy, is phylogenetically widespread but scattered in mammals and birds.

There remain a number of unanswered questions as to how dormancy in endotherms evolved. First, there is considerable debate as to whether daily torpor and hibernation are discrete types of dormancy that emerged independently (Fig. 1A), or if they form a continuum that ranges from no dormancy, to daily torpor, to hibernation (Fig. 1B). On one hand, the discrete states hypothesis is supported by the distributions of most dormancy variables (e.g., minimum body temperature during torpor), which tend to be bimodal, with few species occupying the intermediate space (Ruf & Geiser 2015). On the other hand, the dormancy continuum hypothesis is supported by an interspecific comparison of the “heterothermy index” (a metric of the fluctuations in body temperature over a given time period; Boyles *et al*. 2011), which revealed a continuous distribution (Boyles *et al*. 2013).

**Figure 1:**
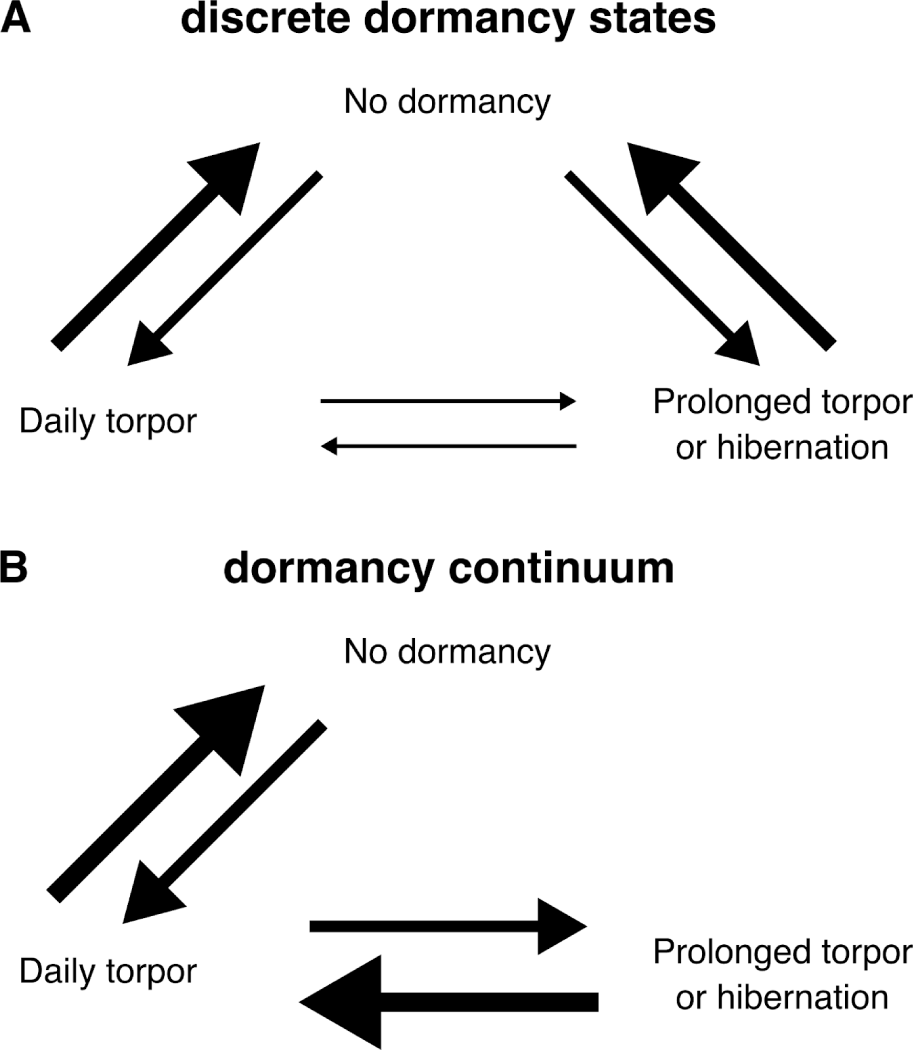
Two alternative hypotheses for the evolutionary transitions among dormancy states in endotherms. Large arrows represent high transition rates, and vice versa. The discrete dormancy states hypothesis (A) posits that daily torpor and hibernation are highly distinct strategies that evolved independently and, therefore, evolutionary transitions between these two states are rare. In contrast, the dormancy continuum hypothesis (B) assumes that daily torpor and hibernation are parts of a continuum and, thus, direct evolutionary shifts between lack of dormancy and hibernation should not be possible.

Second, we also have an incomplete understanding of the physiological and ecological drivers that underlie evolutionary shifts in dormancy. More precisely, it is not known whether, across all endotherms, the ability to enter dormancy is strongly associated with specific ecophysiological variables (e.g., body mass, diet, ambient temperature seasonality), or whether such associations are weak and clade-dependent. If the former is true, it should be possible to predict—to an extent—the propensity for dormancy of species for which such information is currently lacking, based on their physiology and ecological strategies. Identifying strong links between dormancy and ecophysiological variables would also shed light on the costs and benefits of daily torpor and hibernation in different environments, allowing us to more accurately forecast the impacts of global environmental change on species’ populations.

Third, the origins of hibernation remain under debate. In particular, it has been argued that because of the complex physiological underpinnings of hibernation and because hibernation has been observed in monotremes, marsupials, and placentals, their last common ancestor should have been capable of hibernation, with this trait later being lost in many lineages (e.g., see Geiser 1998; Grigg *et al*. 2004; Malan 1996). Furthermore, it has been hypothesised that months-long hibernation may have been key for the survival of mammals after the asteroid impact at the Cretaceous-Palaeogene boundary (e.g., Geiser *et al*. 2017; Lovegrove *et al*. 2014; Nowack *et al*. 2017), which radically restructured ecosystems on a global scale, causing global cooling, wildfires, acid rain, and, ultimately, a mass extinction, including of non-avian dinosaurs (Carvalho *et al*. 2021; Chiarenza *et al*. 2020; Gulick *et al*. 2019; Maruoka 2019). The alternative hypothesis would be that different clades of mammals independently evolved hibernation at various time points, which would suggest that hibernation was not the primary driver of their survival at the Cretaceous-Palaeogene boundary.

To fill the aforementioned knowledge gaps, we compiled the largest dataset of dormancy capabilities to date, covering 1,338 extant mammals and birds, along with 21 physiological and ecological variables potentially associated with dormancy. We then performed a thorough phylogenetic comparative analysis to address three key questions:

1. Are daily torpor and hibernation evolutionarily discrete dormancy states or parts of a continuum?
2. Are there strong associations between dormancy and key ecophysiological variables?
3. Were the early ancestors of modern mammals and birds capable of hibernation or did this trait evolve multiple times in different clades?

## Methods

### Data analysed in the present study

To address our three key questions, we compiled a comprehensive dataset of dormancy capabilities of extant mammals and birds by combining multiple previously published datasets (Geiser 2020, 2021; McNab 2008, 2009; Nowack *et al*. 2020, 2023; Revelo Hernández *et al*. 2023; Ruf & Geiser 2015) and classified each species as belonging to one of three groups: i) no dormancy, ii) daily torpor, and iii) hibernation. This resulted in a dormancy dataset of 737 mammalian and 601 avian species, representing 52 orders according to the NCBI Taxonomy database (Schoch *et al*. 2020).

We additionally collated a dataset of 21 ecophysiological variables for each species, based on literature information where possible: 1) body mass; 2) basal metabolic rate; 3) brain mass; 4) maximum longevity; 5) migration; 6-7) carnivory (including insectivory) and herbivory; 8) fossoriality; 9-12) diurnality, crepuscularity, nocturnality, and cathemerality; 13) aquatic affinity; 14) range size; 15-16) absolute latitude and hemisphere of the centre of the species range; 17-21) mean temperature, temperature seasonality, annual precipitation, precipitation seasonality, and net primary productivity at the centre of the range. Besides extant species, we queried palaeobiological studies to collect additional data for some of our response variables for a few early endotherm ancestors. Descriptions, units, and sources for each variable are available in Supplementary Section S2.

### Estimation of a time-calibrated phylogeny

To obtain a phylogeny of the species in our datasets, we combined the time-calibrated mammalian phylogeny of Álvarez-Carretero *et al*. (2022) and the non-time-calibrated avian phylogeny of Brown *et al*. (2017). For species in our datasets that were not included in these phylogenies, we checked if they were present under a different name. If this was not the case, we manually added these species based on topological information from the Open Tree of Life (Hinchliff *et al*. 2015). We then removed species that were present in the original phylogenies but not in our datasets. The resulting tree was time-calibrated through “congruification” (Eastman *et al*. 2013), which is implemented in the `geiger` R package (v. 2.0.10; Pennell *et al*. 2014). This approach involves identifying nodes in our tree that are compatible with those of time-calibrated reference trees, and transferring age information from the latter to the former. The remaining nodes are calibrated based on penalized likelihood with `treePL` (Smith & O’Meara 2012). As reference trees, we used the Álvarez-Carretero *et al*. (2022) mammalian phylogeny and the phylogeny of the TimeTree database (Kumar *et al*. 2022).

### Phylogenetic comparative analyses

#### Reconstructing the evolutionary pattern of dormancy shifts

To examine how shifts in dormancy occur across the phylogeny, we fitted 12 alternative M*k* models (Lewis 2001) using the `phytools` R package (v. 1.8-4; Revell 2012) that differ in the following three features. First, half of these models allowed evolutionary transitions to occur between any two dormancy states (in line with the discrete dormancy states hypothesis; Fig. 1A), whereas, in the remaining half, direct transitions between no dormancy and hibernation were not allowed (in line with the dormancy continuum hypothesis; Fig. 1B). Second, models differed in their patterns of transition rates between pairs of dormancy states, assuming a) a common rate for all possible transitions, or b) a common rate for forward and backward transitions (e.g., torpor to hibernation and hibernation to torpor), or c) a different rate for each possible transition. Third, for half of the models, we simultaneously estimated both the transition rate(s) and the value of Pagel’s λ (Pagel 1999). This allowed us to account for the possibility that dormancy may not exhibit a strong phylogenetic signal, i.e., closely related species may not necessarily be considerably more similar in their dormancy capabilities than randomly selected species. After obtaining all model fits, we calculated the Akaike weight (Akaike 1974) for each candidate model and performed model averaging to obtain the final transition rates.

#### Estimation of associations between dormancy and ecophysiological variables

To investigate the links among dormancy and key physiological and ecological factors, we fitted a multi-response generalised linear mixed model using the `MCMCglmm` R package (v. 2.34; Hadfield 2010, 2015). This computationally intensive model (requiring ∼180 CPU hours per Markov chain; Supplementary Section S3.4) had 22 response variables, corresponding to dormancy and our 21 ecophysiological variables described earlier. Thus, the model simultaneously predicted all response variables along with their variance-covariance matrix, enabling us to estimate pairwise correlations among responses. In such a model, each response variable needs to conform to one of the distributions implemented in `MCMCglmm` (e.g., Gaussian, Poisson). To this end, we transformed continuous variables towards approximate normality when that was necessary. More precisely, we applied i) a natural logarithm transformation to body mass, basal metabolic rate, brain mass, maximum longevity, and annual precipitation, ii) a square root transformation to absolute latitude and temperature seasonality, iii) a cube root transformation to net primary productivity, iv) a fourth root transformation to precipitation seasonality, and v) a fifth root transformation to range size.

We modelled categorical variables as threshold variables. The threshold model originates from the field of quantitative genetics (Wright 1934) and was later applied in a phylogenetic comparative context by Felsenstein (2005). It assumes that a categorical variable is governed by an underlying unobserved continuous trait called the “liability”, and that shifts from one state to another occur when the liability moves past a particular threshold value. The threshold model can accommodate both binary and multistate categorical variables, provided that the states follow a particular order (e.g., we coded dormancy as three ordered states of no dormancy, daily torpor, and hibernation). In the latter case, the model will also identify the location of the additional threshold(s) in liability space. We, therefore, applied the threshold approach—as implemented in `MCMCglmm`—to dormancy and all other categorical variables in our dataset, ordering their states as shown in Supplementary Table S1.

In addition to data for extant species, we were able to include data for ancestral nodes (Supplementary Section S2.2) in the model. A special case was the last common ancestor of Amniota, which includes mammals and birds. Its habitat was likely coal swamps (Clack 2012), suggesting either moderate or low aquatic affinity, whereas its body mass was estimated by Brocklehurst et al. (2022) at less than 1 kg. To incorporate this uncertainty into our parameter estimates, we specified 6 alternative models using `MCMCglmm`, with all possible combinations of a) a low or moderate aquatic affinity, and b) a body mass of 2, 500, or 1,000 g, and combined their posterior distributions (Supplementary Section S3.4).

We included no explanatory variables in the models, other than a separate intercept per response variable, on which we specified a phylogenetic random effect to account for the evolutionary relationships across species. We summarised the posterior distributions of model parameters (e.g., correlations between response variables, thresholds) by calculating the median of each distribution and the 95% Highest Posterior Density (HPD) interval. Given that body mass correlated strongly with metabolic rate, brain mass, and maximum longevity, as expected, we removed the contribution of body mass from each of these three traits (Supplementary Section S3.5) and fitted the models again using the mass-corrected traits. Further details on the specification of these models are available in Supplementary Section S3.

#### Comparison of dormancy-capable species

To understand how dormancy-capable species differ from each other, we compared them based on the 21 ecophysiological variables in our study. To this end, we first extracted the posterior distributions of the 21 variables per species from the `MCMCglmm` fits. For each posterior distribution, we compiled a table of the median and the bounds of the 95% HPD interval. By doing so, we were able to a) include species with missing values in one or more variables (Supplementary Section S3.1) and b) explicitly account for the uncertainty around each estimate, which may depend on the pattern of data missingness and the position of the species in the phylogeny. Next, we applied a phylogenetic principal components analysis (pPCA) to the aforementioned table, accounting for the strength of the phylogenetic signal in the data (the λ parameter), as implemented in the `phytools` R package.

We then examined how daily torpor and hibernation are distributed across the first four phylogenetic principal components. Furthermore, to investigate whether there are multiple subtypes of daily torpor and hibernation that are associated with distinct areas of the ecophysiological space (i.e., distinct niches), we applied Gaussian mixture model clustering to pPCA scores using the `mclust` R package (v. 6.0.0; Scrucca *et al*. 2016), setting the number of clusters between 1 and 5. The optimal mixture model was automatically determined using the Bayesian Information Criterion (Schwarz 1978). We additionally examined the distribution of three dormancy descriptors introduced by Nowack *et al*. (2023) across the first four phylogenetic principal components (Supplementary Section S6.1).

#### Ancestral state reconstruction of dormancy and robustness analysis

To infer if dormancy was present in early ancestors of modern mammals and birds, we reconstructed the ancestral dormancy states of major avian and mammalian clades according to both the M*k* and `MCMCglmm` fits. For the latter model, we extracted the posterior dormancy probabilities that were already calculated for all internal nodes of the phylogeny during the fitting process. For the M*k* model, we conducted 10,000 stochastic character mapping simulations (Bollback 2006; Huelsenbeck *et al*. 2003) using the `make.simmap` function of the `phytools` R package. In these simulations, we set the transition rate matrix and the dormancy probabilities at the root of the tree to the previously obtained model-averaged estimates.

Given that a) placental mammals were the group with the highest number of dormancy shifts in our dataset and b) our knowledge of which species are capable of dormancy remains incomplete, we further investigated the robustness of the dormancy probabilities for the last common ancestor of modern placental mammals. To this end, we assumed that a certain number of placental mammals, currently not known to be dormancy-capable, are in fact dormancy-capable, and randomly switched 20, 85, or 200 of them from no dormancy to either daily torpor or hibernation. These correspond to 5.08%, 21.57%, and 50.76% of the dormancy-incapable placental mammals in our dataset. We performed this process 5 times, fitted the M*k* and `MCMCglmm` models to all 15 resulting datasets, and re-estimated the dormancy probabilities of Placentalia as described earlier.

All computational analyses described in this study were executed on compute nodes equipped with AMD EPYC 7702 CPU cores for a cumulative runtime of ∼101,600 CPU hours (see also Supplementary Section S3.4).

## Results

### Evolutionary transitions among dormancy states

To address whether daily torpor and hibernation are discrete dormancy states or parts of a continuum (Fig. 1), we reconstructed the pattern of evolutionary transitions among these three states based on fits of the M*k* model. We found strong support for frequent evolutionary transitions between no dormancy and daily torpor, and between daily torpor and hibernation, but direct transitions between no dormancy and hibernation were nearly nonexistent (Fig. 2B). Considering the possibility that the few observed apparent direct transitions (Supplementary Section S1) may involve a transient and unobserved daily torpor state, this result provides strong support for the continuum hypothesis. It also justifies our treatment of dormancy as a threshold trait in our multi-response generalised linear mixed models. In these models, median estimates of dormancy liability for the 1,338 species covered nearly the entire continuum (Fig. 2B,C). For example, liabilities of species capable of daily torpor ranged from relatively low (just above the no-dormancy threshold) to intermediate to relatively high (just below the hibernation threshold).

**Figure 2.**
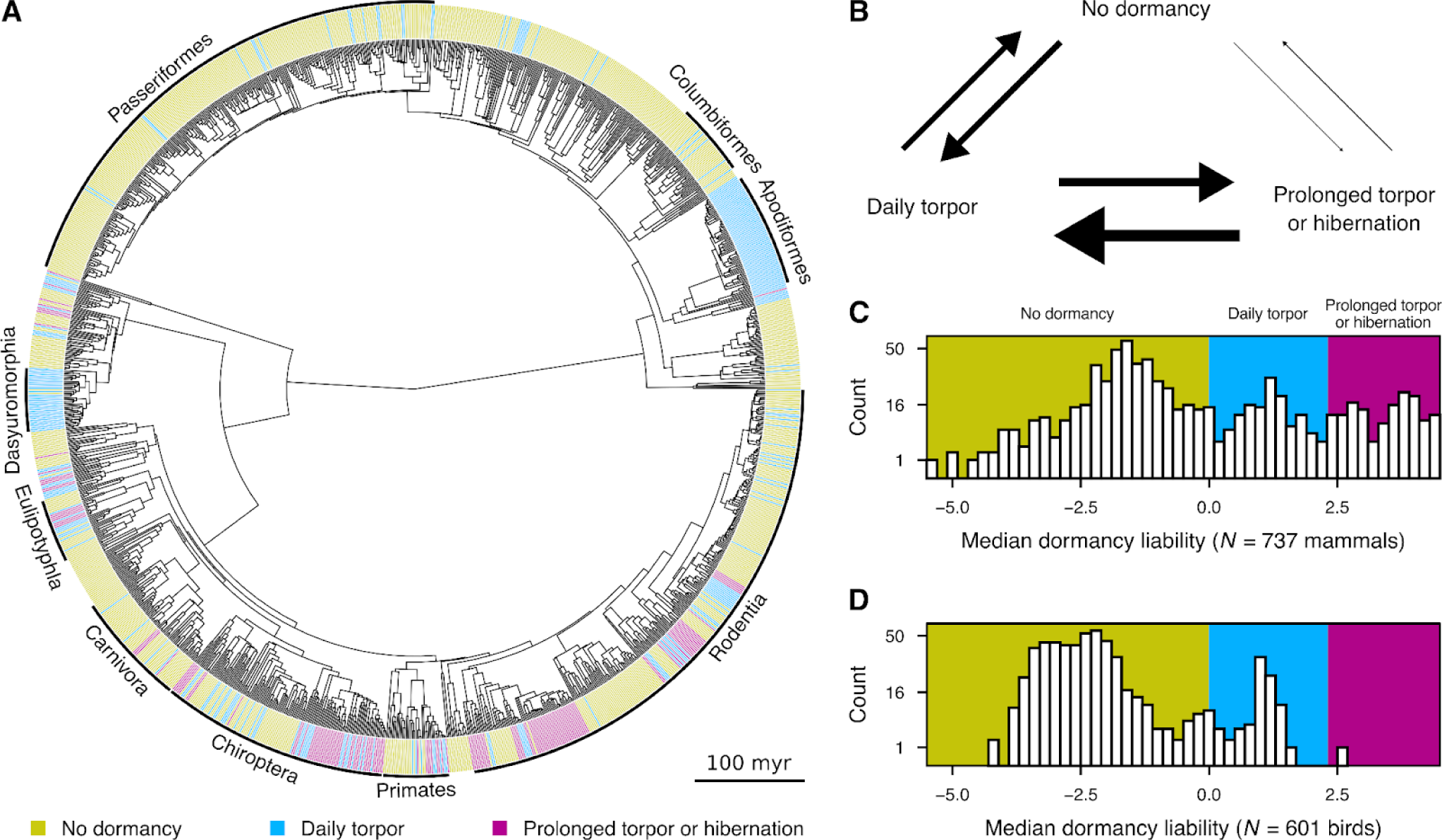
(on the previous page): Distribution of dormancy across the phylogeny of mammals and birds and evolutionary patterns thereof. (A) The most species-rich orders in our dataset are explicitly shown. (B) The pattern of evolutionary shifts among dormancy states, as reconstructed with variants of the M*k* model. (C-D) Median posterior dormancy liabilities for all extant mammals (C) and birds (D) in our dataset, as estimated with `MCMCglmm` fits (see Methods). Thresholds are denoted by a shift in background colour. Note that values along the vertical axes do not increase linearly so that both small and large counts are distinguishable.

### Associations of dormancy with ecophysiological factors

We next examined the associations between dormancy and the 21 ecophysiological variables that we collected. Across all 1,338 extant mammalian and avian species in our dataset, we were able to detect systematic correlations with dormancy (i.e., with a 95% HPD interval that does not include zero) for 16 of the 21 variables (Fig. 3). For those, median posterior correlation estimates ranged from -0.33 to -0.13 (negative associations) and from 0.12 to 0.32 (positive associations). The variables that correlated most strongly and positively with the ability to enter dormancy were temperature seasonality and absolute latitude at the middle of each species’ range, nocturnality, and carnivory. In contrast, dormancy was most strongly negatively correlated with net primary productivity, annual precipitation, and mean temperature at the range centre, as well as with body mass and cathemerality.

**Figure 3:**
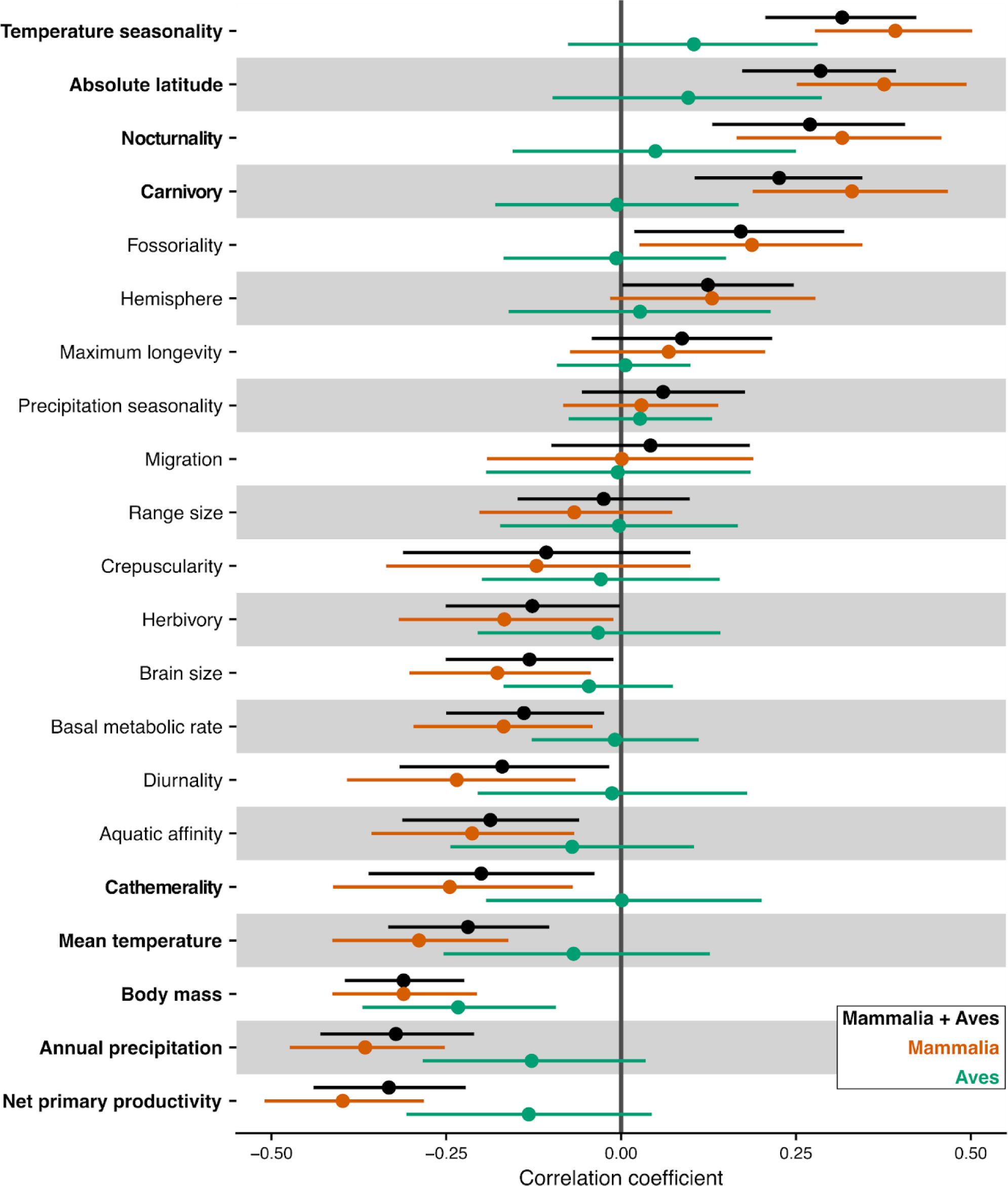
Correlations between dormancy and ecophysiological variables. Circles stand for the median posterior estimate of each correlation coefficient, whereas horizontal lines represent its 95% HPD interval. Systematic correlations are considered as those with a 95% HPD interval that does not include zero. Variables shown in bold have an absolute median correlation estimate of at least 0.2 across the entire dataset (Mammalia + Aves). Basal metabolic rate, brain mass, and maximum longevity were corrected for body mass (Supplementary Section S3.4).

Next, we investigated associations between dormancy and ecophysiological variables, separately for mammals and for birds. Including only mammals in the correlations analysis produced qualitatively identical results for most variables. Specifically, correlations obtained with the entire dataset generally increased in strength when only mammals were included. The main exception was the correlation with hemisphere which was marginally supported across the entire dataset and not supported for the mammalian subset. In contrast, including only birds in the analysis resulted in much weaker correlations with dormancy for all 21 variables. In particular, the correlation with body mass was the only systematic correlation for the avian subset of the data.

### Variation among dormancy-capable species

A phylogenetic PCA (pPCA) of the 21 ecophysiological variables across only dormancy-capable species revealed extensive overlap between daily torpor and hibernation in the parameter space (Fig. 4). Clustering of the resulting pPCA scores did not support the existence of multiple subtypes of daily torpor and hibernation, tied to specific physiological, ecological, or climatic factors. Instead, variation among dormancy-capable species was strongly taxonomically structured (Fig. 5). Analysing mammals separately and examining the distribution of the three dormancy descriptors of Nowack *et al*. (2023) across the first four pPCA principal components led to similar conclusions (Supplementary Section S6).

**Figure 4:**
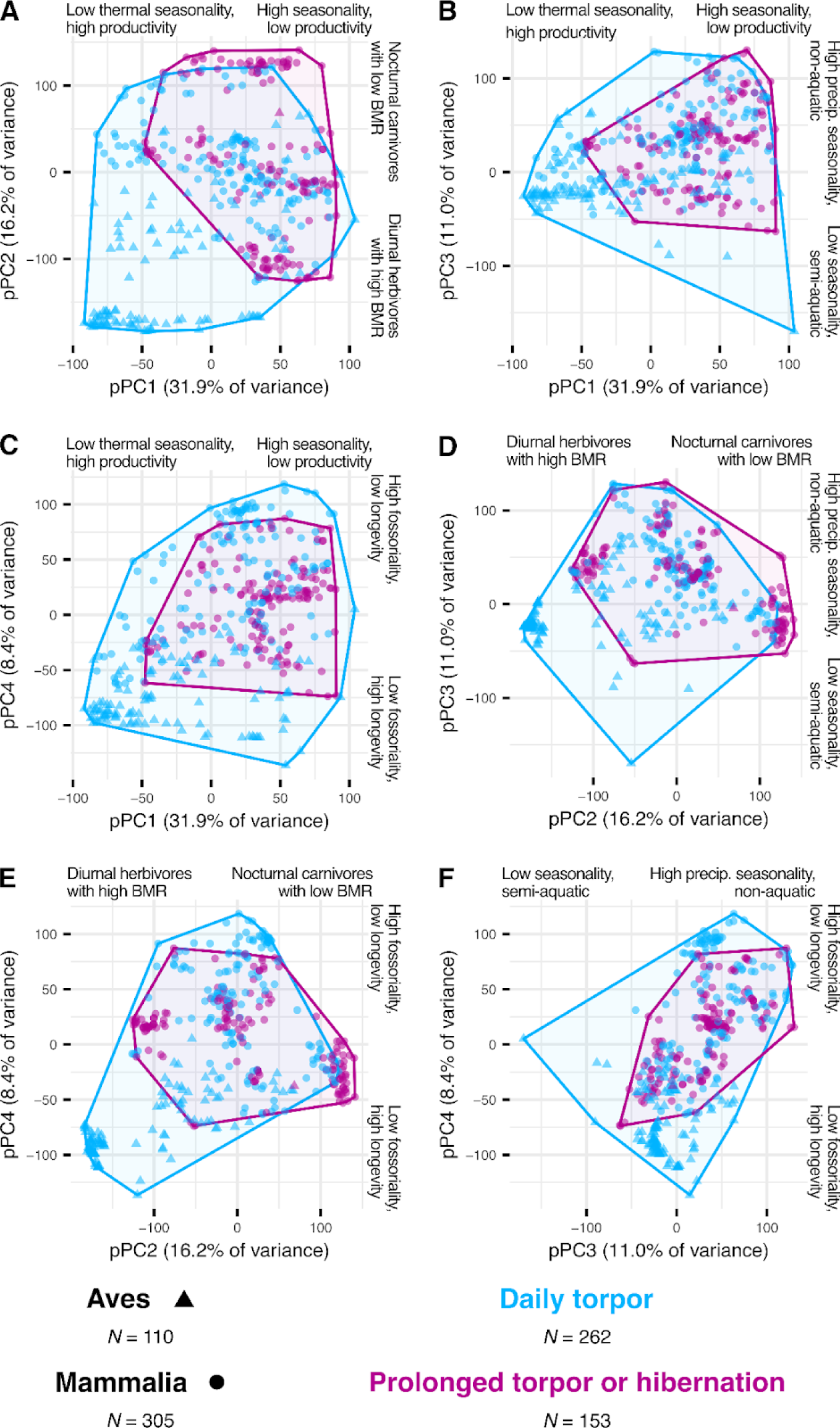
Projection of daily torpor (blue) and hibernation (purple) onto the first four principal components of a phylogenetic PCA of 21 ecophysiological variables for dormancy-capable species only (see Methods). For each principal component, the variables that most strongly correlate with it are explicitly listed.

**Figure 5:**
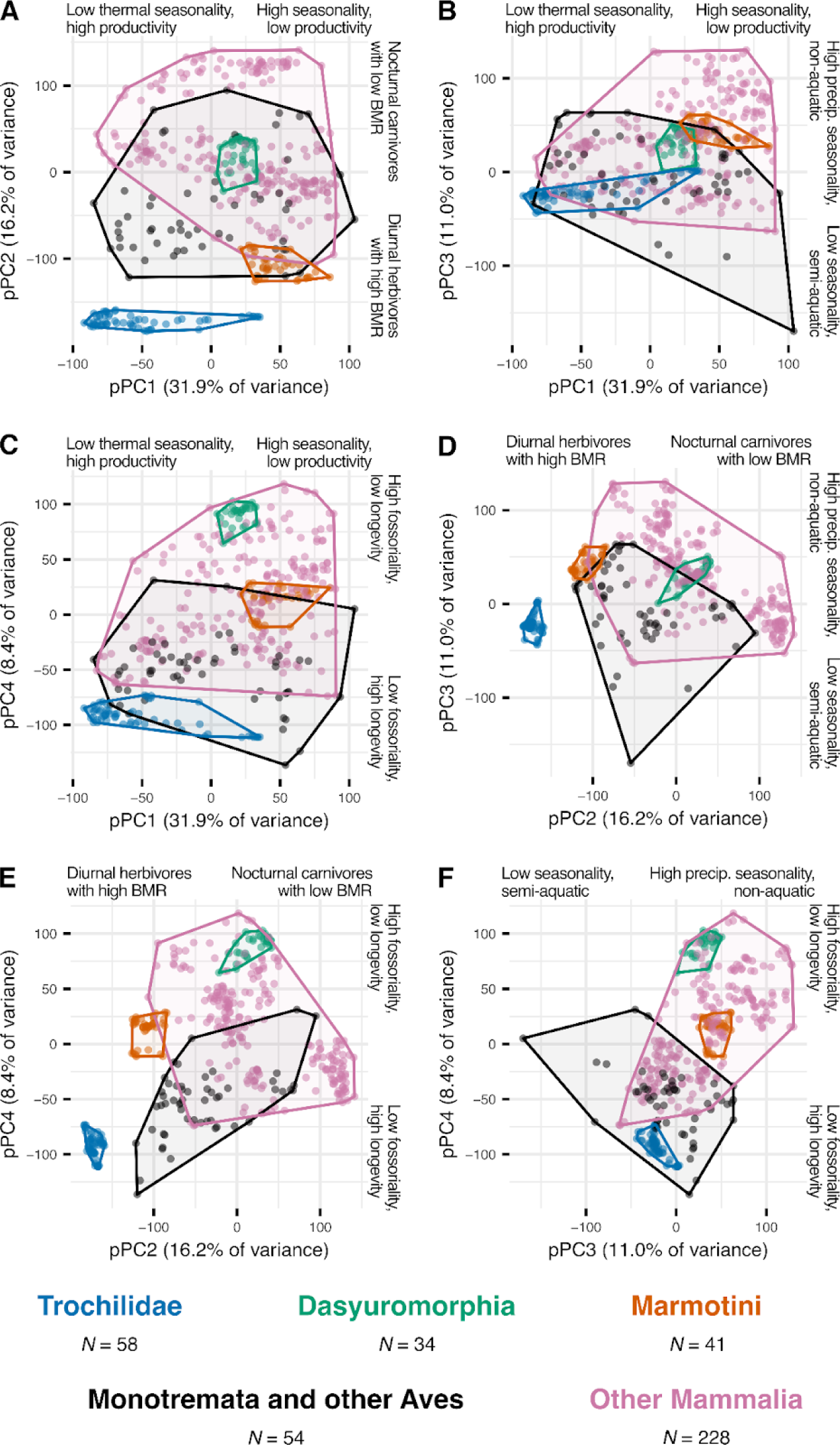
Clustering of dormancy-capable species based on their distribution throughout the ecophysiological parameter space, as inferred from the phylogenetic PCA. Panels A-F show all pairwise combinations of the first four phylogenetic principal components. Note that three of the five clusters are monophyletic.

### Origins of hibernation in mammals and birds

We next inferred whether the early ancestors of mammals and early ancestors of birds were capable of hibernation. In all deep nodes that we examined, we estimated a low probability for hibernation, according to both M*k* and `MCMCglmm` fits. For the last common ancestor of extant placental mammals (LCAP), we additionally tested the robustness of this estimate to assuming that our knowledge of dormancy-capable species is incomplete by randomly switching 20, 85, or 200 of the dormancy-incapable placental mammals to either daily torpor or hibernation. This test showed that our finding of a low probability for hibernation for the LCAP was almost always robust (Fig. 6C,D). We only obtained a high probability of hibernation for two of the five tests when switching the dormancy capabilities of 200 species. However, in these two tests, the M*k* model assigned a non-negligible weight to direct evolutionary transitions between no dormancy and hibernation (Fig. 6C,E). In other words, even after switching >50% of the dormancy-incapable placental mammals, a high probability of hibernation in the LCAP is only possible if daily torpor and hibernation are discrete evolutionary states, which violates the strongly supported dormancy continuum hypothesis. This also explains why MCMCglmm fits—which necessarily impose a continuum in dormancy—never estimated a high probability of hibernation in the LCAP, even when we modified the dormancy capabilities of 200 species (Fig. 6D).

**Figure 6:**
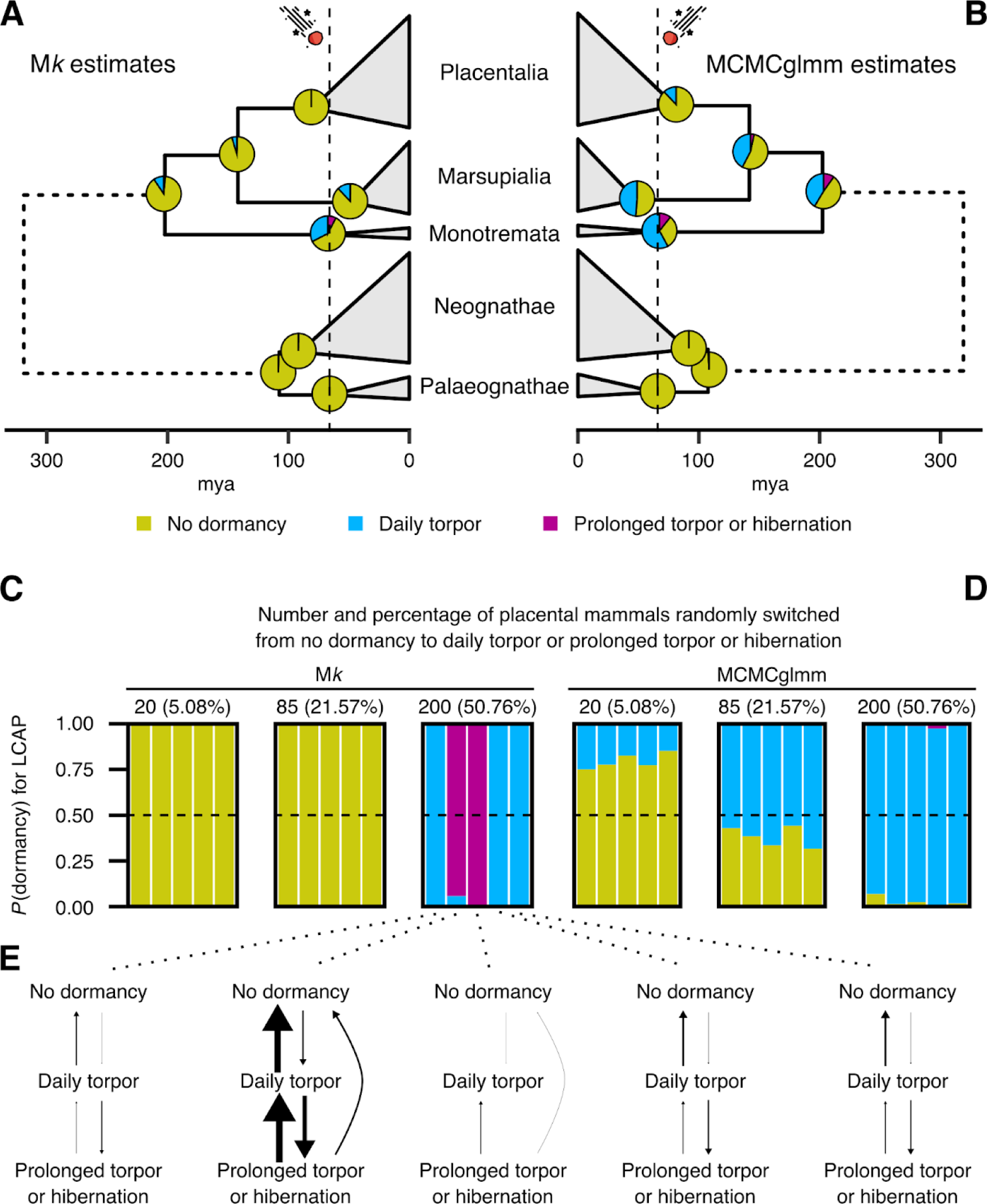
Ancestral state reconstruction of dormancy for selected deep nodes of our phylogeny. (A-B) Dormancy probabilities according to M*k* and MCMCglmm fits. The dashed vertical line at 66 mya indicates the Cretaceous-Palaeogene boundary. (C-D) Probabilities of the three dormancy states for the last common ancestor of Placentalia (LCAP), according to M*k* and MCMCglmm fits after randomly switching 20, 85, or 200 dormancy-incapable placental mammals to dormancy-capable, replicated 5 times. (E) Reconstructed evolutionary transitions among dormancy states, as inferred with variants of the M*k* model, after switching 200 placental mammals from dormancy-incapable to either daily torpor or hibernation.

## Discussion

In this study, we compiled a comprehensive dataset of dormancy capabilities of 1,338 extant mammals and birds, along with 21 ecophysiological variables for each species and for a few ancestral taxa based on palaeobiological studies. By applying a series of phylogenetic comparative methods to the data, we addressed three key questions regarding the evolution of dormancy as a key survival strategy for many mammals and birds.

Our first key question was whether daily torpor and hibernation are evolutionarily distinct states or rather form a dormancy continuum (Fig. 1). Our results strongly support the dormancy continuum hypothesis, as a) direct transitions between no dormancy and hibernation were very rarely observed (Fig. 2B), and b) daily heterotherms and hibernators overlapped remarkably in the ecophysiological space (Fig. 4). Direct transitions between no dormancy and hibernation were detected in only 11 branches of our phylogeny, distributed across various groups, including in the Sciuromorpha, Arctoidea, and Xenarthra (Supplementary Section S1). Some direct transitions may be artifactual, arising from errors in tree topology or branch lengths, errors in dormancy classification (e.g., a daily heterotherm being classified as incapable of dormancy), or strong increases in the evolutionary rate of dormancy in some lineages, possibly driven by adaptation to novel environments, that could have resulted in a transient but unobserved daily torpor state on the path to hibernation. Differentiating among these possible explanations for a given direct transition would require further fieldwork, experiments, and computational investigation, and can be the focus of future studies. An interesting example is the dormancy-incapable white-tailed antelope squirrel (*Ammospermophilus leucurus*), which accounts for one of the three direct transitions between no dormancy and hibernation in the Sciuromorpha suborder (Supplementary Fig. S1). *A. leucurus* is nested within a clade of 33 hibernators, but the species has been well-studied both in its natural environment and experimentally, and there is currently no evidence that it can enter controlled dormancy (Karasov 1983; Refinetti & Kenagy 2018, 2023). Thus, overall, our analyses show that dormancy largely evolves along a continuum, with very few possible exceptions.

The second question that we addressed was whether dormancy in endotherms is strongly linked to specific ecophysiological factors, regardless of taxonomic group. To this end, we treated dormancy as a threshold trait—justified by the aforementioned support for a dormancy continuum—and simultaneously estimated the correlation structure among dormancy and 21 ecophysiological variables, accounting for phylogeny. Across all endotherms in the present study, several variables were statistically correlated with dormancy (Fig. 3). Specifically, we found that dormancy is linked with living in a low-productivity environment with high thermal seasonality, nocturnal activity patterns, an at least partly carnivorous diet (including insectivory), and a small body mass. However, we should note that these correlations are overall weak, with median posterior correlation estimates not being higher than 0.32 or lower than -0.33.

The detected associations between dormancy and environmental conditions are in line with the expectation that dormancy should be most beneficial in environments with seasonal variation in temperature and food availability, and less so in thermally stable, highly productive environments. Nevertheless, such associations may also partly reflect the fact that most research effort has been focused on species living in seasonal environments (Levesque *et al*. 2023; Nowack *et al*. 2020), possibly resulting in an underestimation of the dormancy capabilities of species at low latitudes.

Our finding of a link between dormancy and nocturnal activity patterns is consistent with nocturnality being more energetically demanding than diurnality, given that nocturnal species are active under lower ambient temperatures than their diurnal counterparts (Levy *et al*. 2012; van der Vinne *et al*. 2015, 2019). When energetically challenged, some nocturnal species tend to shift to diurnality (e.g., see Hut *et al*. 2012; van der Vinne *et al*. 2019; Weyer *et al*. 2020) and, therefore, undergoing dormancy would be another highly beneficial strategy. Nocturnal species also have the advantage of being able to rewarm passively with rising ambient temperatures which lessens the costs of rewarming from torpor (Lovegrove *et al*. 1999).

The positive association between dormancy and carnivory is likely also driven in part by climate, given that the abundances of many insect species—that serve as prey for insectivorous endotherms—can strongly vary over time, as a function of factors such as temperature and precipitation (e.g., see Bowles *et al*. 2002; Guédot *et al*. 2018; Kishimoto-Yamada & Itioka 2015). It is possible that the different macronutrient compositions of carnivorous and herbivorous diets may play an additional role in the evolution of dormancy, but research on this topic is lacking.

At the physiological level, a negative correlation of dormancy with body mass partly represents a biophysical constraint, given that the maximum rates of cooling and rewarming are inversely proportional to body mass (Geiser 2021; Geiser & Baudinette 1990; McKechnie & Wolf 2004). This suggests that above a certain body mass threshold, the energy expenditure at the beginning and end of dormancy would exceed the amount of energy conserved during dormancy, making it an unfavourable strategy. A key exception to this dormancy-size “rule” are hibernating bears. While their metabolic rate is greatly reduced during hibernation, their body temperature falls to only around 30°C (Evans *et al*. 2016; Hissa 1997; Tøien *et al*. 2011), in contrast to small hibernators such as the Arctic ground squirrel (*Urocitellus parryii*) and the hazel dormouse (*Muscardinus avellanarius*) whose body temperature can become as low as -2.9°C (Barnes 1989; Pretzlaff & Dausmann 2012). This allows bears to enter and exit dormancy relatively quickly, enabling hibernation despite their high body mass.

Including only mammals in the correlations analysis yielded similar results, with most correlations increasing in strength, whereas, across birds, the only systematic correlation for dormancy was a weak negative correlation with body mass. The absence of multiple systematic correlations in birds may be explained by multiple factors. First, compared to mammals, birds have a much sparser distribution of dormancy which—with the exception of hummingbirds—tends to be more scattered across avian clades (Fig. 2A). This would necessarily lower the signal-to-noise ratio across birds compared to mammals, hindering the detection of systematic associations. Second, the increased mobility of flying species which facilitates migration likely contributes to the apparent decoupling of dormancy from environmental variables in birds. Many birds employ seasonal migration to avoid adverse environmental conditions, a strategy that is sometimes combined with dormancy to reduce energy expenditure at stopovers, decreasing the time required for refuelling (Carpenter & Hixon 1988; Clerc & McGuire 2021; Eberts *et al*. 2021; McGuire *et al*. 2023; Wojciechowski & Pinshow 2009). It is also worth pointing out that the common poorwill (*Phalaenoptilus nuttallii)*, the only bird known to hibernate, also migrates in a seasonal manner (Csada & Brigham 1994; Woods & Brigham 2004). Such behaviours would explain why the climatic conditions at the centre of the range are poor predictors of the dormancy capabilities of bird species. In contrast, across mammals, climatic variables exhibit much higher predictive power for dormancy, given that long-distance migration is mainly limited to large-bodied species (that are typically dormancy-incapable) and bats (in which dormancy-assisted migration has been observed; Supplementary Fig. S12; Baloun & Guglielmo 2019; McGuire *et al*. 2014, 2023; Webber & McGuire 2022).

Our last key question was whether the early ancestors of modern endotherms were capable of hibernation which was later lost by most lineages, or whether multiple lineages evolved hibernation independently. Our results support the latter scenario given that, for the deepest nodes in our phylogeny, hibernation was the least likely state according to both M*k* and `MCMCglmm` fits (Fig. 6A,B). Interestingly, this includes nodes both before and after the Cretaceous-Palaeogene boundary, suggesting that hibernation may not have been a key ability for the survival of certain endothermic lineages during that mass extinction event. Overall, our results suggest that early mammalian and avian ancestors did not hibernate, with this trait emerging independently in many lineages of endotherms, enabling them to occupy a wide diversity of niches.

A scenario of repeated independent gains (and losses) of hibernation and daily torpor may appear counterintuitive given the complex physiological processes that are responsible for entering, maintaining, and exiting dormancy (e.g., see Fu *et al*. 2020; Junkins *et al*. 2022; Mohr *et al*. 2020). Nevertheless, it can explain the remarkable variation in the physiologies, ecological strategies, and dormancy characteristics present among dormancy-capable endotherms (Figs. 3-5, Supplementary Figs. S13-S16). Such a scenario is also consistent with the reconstructed evolutionary transitions among dormancy states (Fig. 2B), given that shifts a) from no dormancy to daily torpor and b) from daily torpor to hibernation are relatively frequent (i.e., the corresponding arrows have a non-negligible size). It is also worth mentioning that many other complex traits have evolved convergently multiple times, including vision (Nilsson 2021), flight (Alexander 2015), extreme longevity (Yu *et al*. 2021), and the return to a fully aquatic lifestyle (Houssaye & Fish 2016).

In conclusion, our study shows that daily torpor and hibernation are not evolutionarily distinct dormancy states and that dormancy likely emerged independently multiple times across endotherms and in different ways. As a result, knowledge of the patterns and molecular underpinnings of dormancy of a given species may not be directly transferable to other, evolutionarily distant species, even if they occupy similar niches. Thus, more attention should be paid to thoroughly characterising the dormancy capabilities and patterns—at the molecular, physiological, or ecological level—of species representing diverse clades. This will both deepen our understanding of the similarities and differences in dormancy across narrow and broad taxonomic groups, and improve our ability to forecast the responses of dormancy-capable endotherms to future environmental challenges.

## Acknowledgements

We thank the members of the Hiller lab for useful discussions and feedback, and Christoph Sinai for technical support. DGK was supported by an EMBO Postdoctoral Fellowship (ALTF 1089-2021). This work was also supported by the LOEWE-Centre for Translational Biodiversity Genomics (TBG), funded by the Hessen State Ministry of Higher Education, Research and the Arts (HMWK).

