## Supporting Information for "Numerous independent gains of torpor and hibernation across endotherms, linked with adaptation to diverse environments"

#### Contents

**S1. Direct shifts between no dormancy and hibernation according to the M*k* model 2**

**S2. Ecophysiological data used in this study 6**

S2.1. Sources of ecophysiological data for extant species 6

S2.2. Sources of ecophysiological data for internal tree nodes 8

S2.2.1. Body mass 8

S2.2.2. Brain mass 8

S2.2.3. Basal metabolic rate 8

S2.2.4. Carnivory and herbivory 8

S2.2.5. Aquatic affinity 8

**S3. Additional MCMCglmm specification details 9**

S3.1. Continuous imputation of missing values in response variables 9

S3.2. Variance-covariance matrices of response variables 9

S3.3. Priors 9

S3.4. Number of generations and posterior diagnostics 9

S3.5. Body mass-correction of metabolic rate, brain mass, and maximum longevity 9

**S4. Comparisons of the predictions of MCMCglmm fits with data 10**

**S5. The relationship between migration and body mass in endotherms 12**

**S6. Further analyses of the distribution of dormancy along the ecophysiological parameter space 12**

S6.1. Projection of dormancy descriptors onto the ecophysiological parameter space 12

S6.2. Phylogenetic PCA across only mammal species 16

**References 17**

**1.** LOEWE Centre for Translational Biodiversity Genomics, Senckenberganlage 25, 60325 Frankfurt am Main, Germany
**2.** Senckenberg Research Institute, Senckenberganlage 25, 60325 Frankfurt am Main, Germany
**3.** School of Biology and Ecology, University of Maine, Orono, ME, 04469, USA
**4.** Institute of Cell Biology and Neuroscience, Faculty of Biosciences, Goethe University Frankfurt, Max-von-Laue-Str. 9, 60438 Frankfurt am Main, Germany

#### S1. Direct shifts between no dormancy and hibernation according to the M*k* model

To identify branches where direct shifts between no dormancy and hibernation occurred, we first performed 10,000 stochastic character mapping simulations using the M*k* model, as described in the main text. We then searched for branches where a direct shift occurred in at least half of the simulations. The total number of such branches was 11 (Supporting Figs. S1-S8).

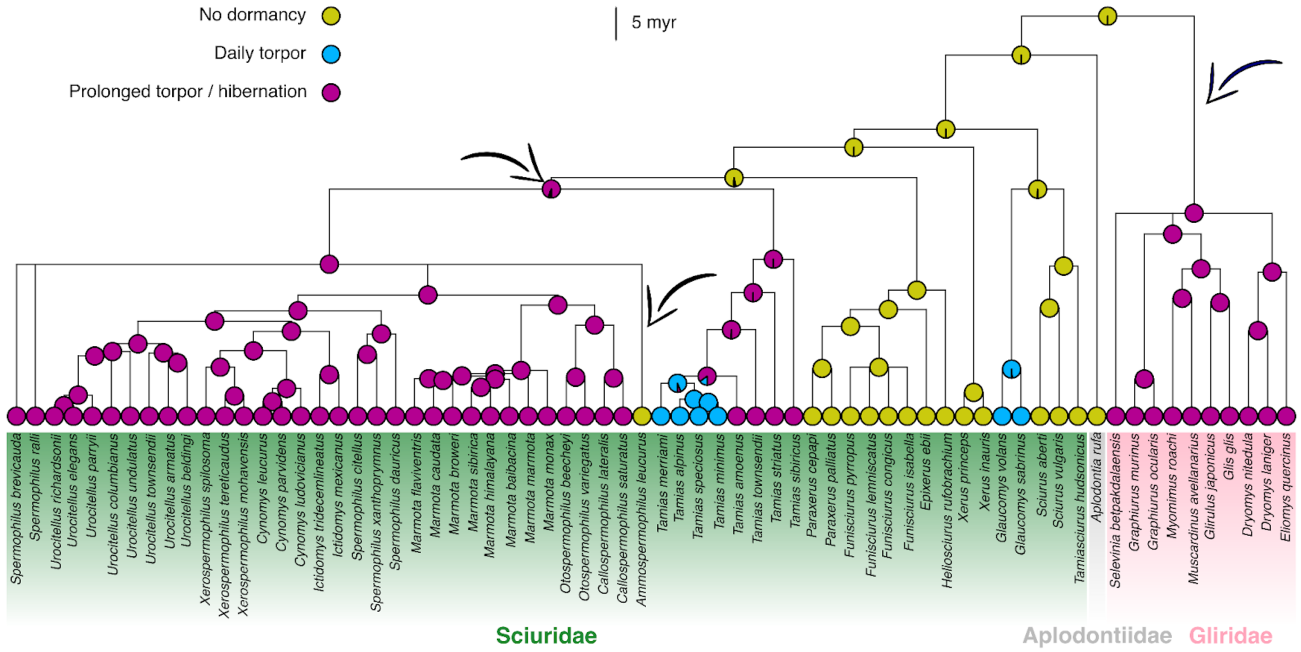


**Figure S1:** Three direct shifts between no dormancy and hibernation, denoted by arrows, across the Sciuromorpha suborder. Families are shown in different colours.

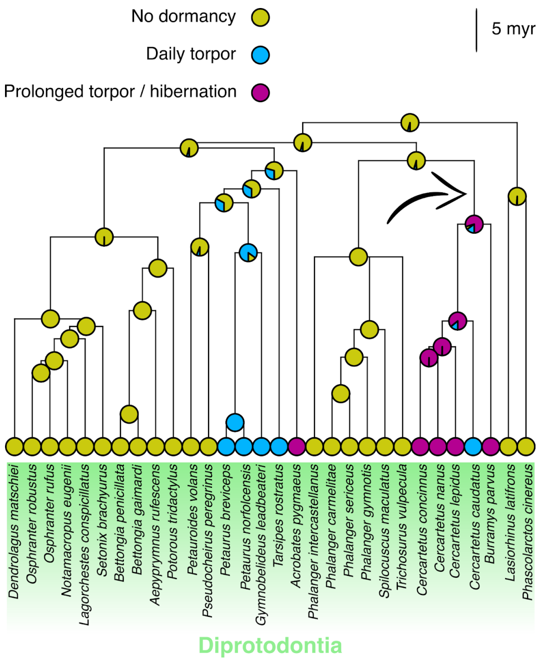


**Figure S2:** A direct shift between no dormancy and hibernation, denoted by an arrow, across the Diprotodontia order.

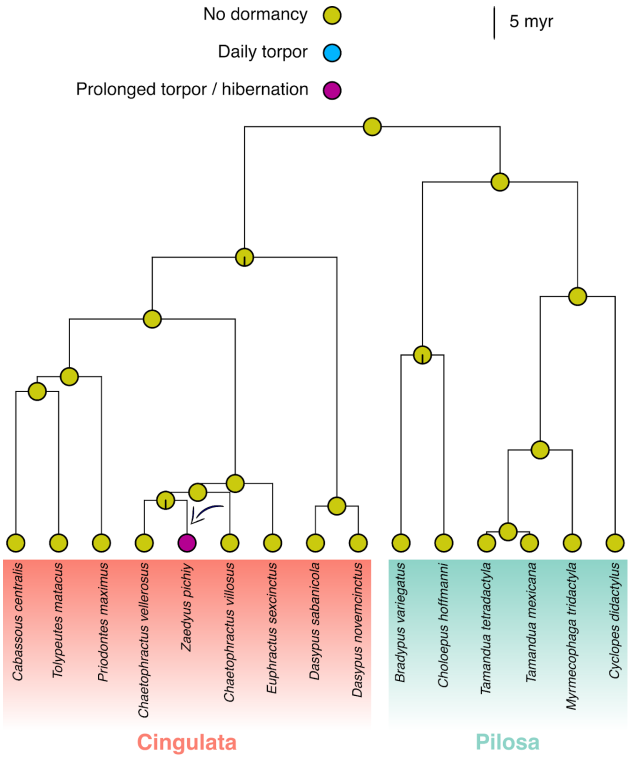


**Figure S3:** A direct shift between no dormancy and hibernation, denoted by an arrow, across the Xenarthra superorder. Orders are shown in different colours.

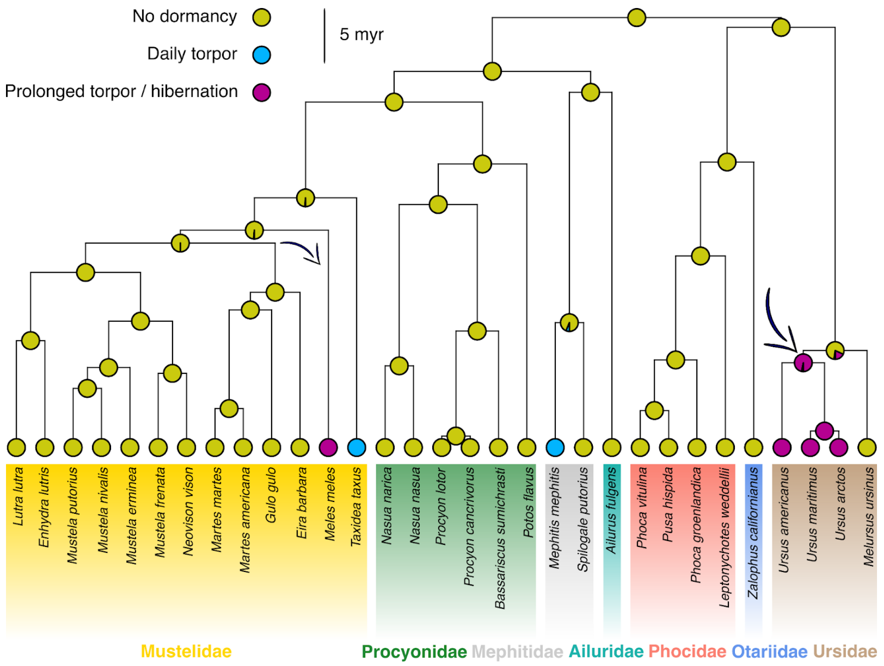


**Figure S4:** Two direct shifts between no dormancy and hibernation, denoted by arrows, across the Arctoidea infraorder. Families are shown in different colours.

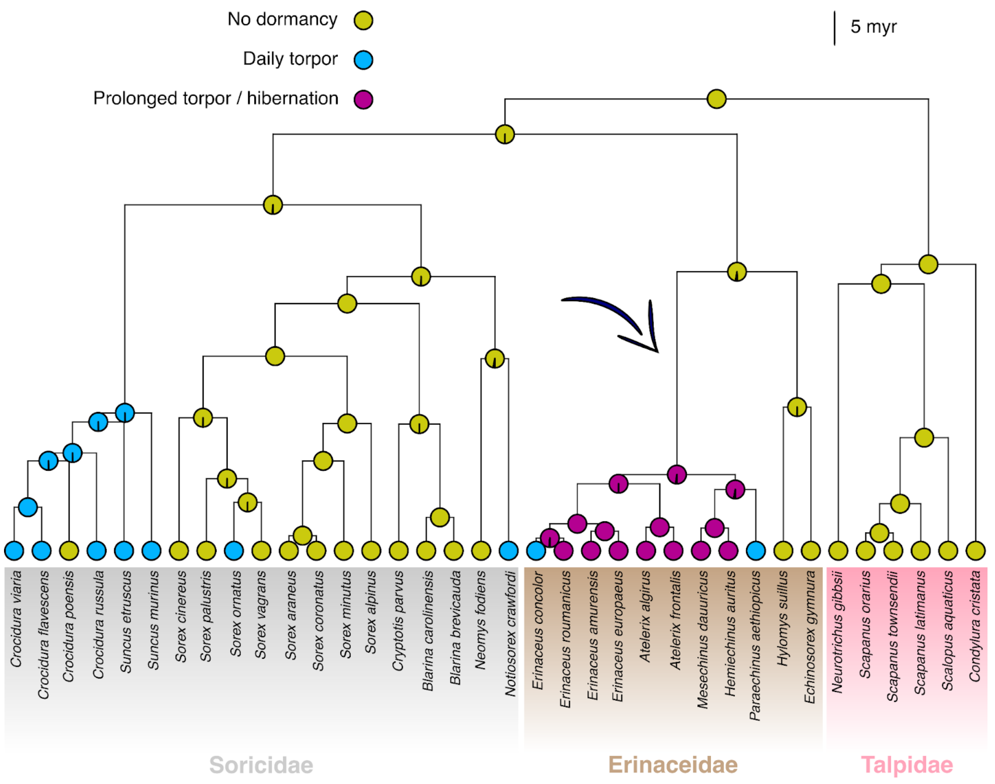


**Figure S5:** A direct shift between no dormancy and hibernation, denoted by an arrow, across the Eulipotyphla order. Families are shown in different colours.

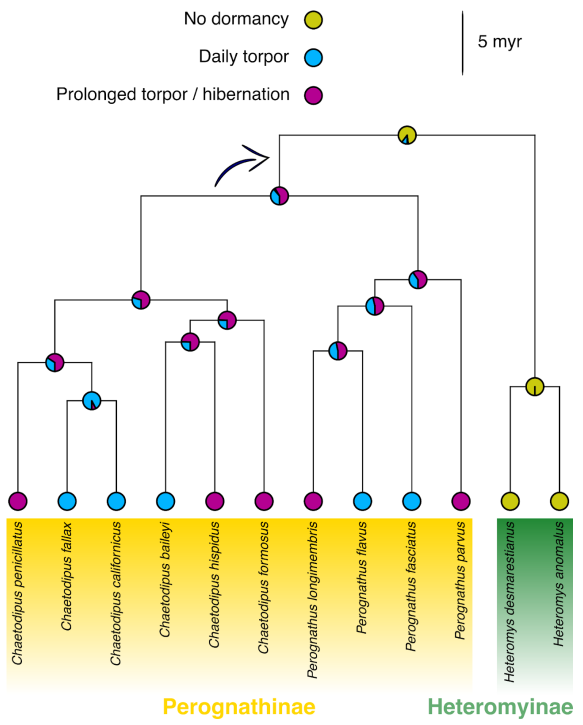


**Figure S6:** A direct shift between no dormancy and hibernation, denoted by an arrow, across the Heteromyidae family. Subfamilies are shown in different colours.

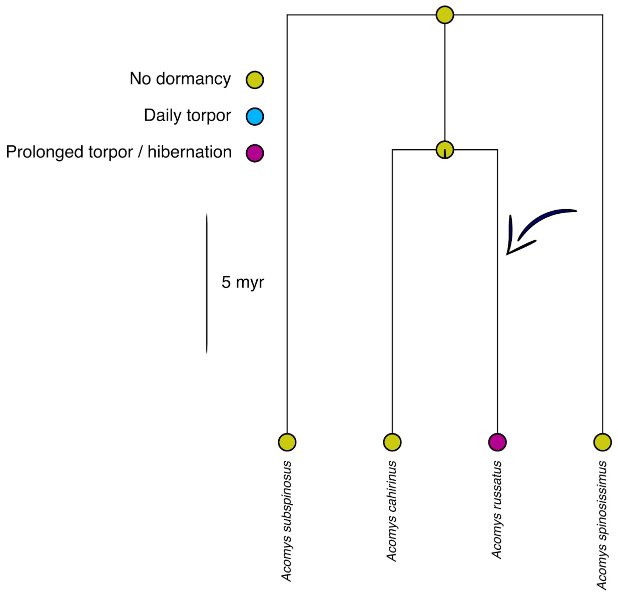


**Figure S7:** A direct shift between no dormancy and hibernation, denoted by an arrow, across the *Acomys* genus.

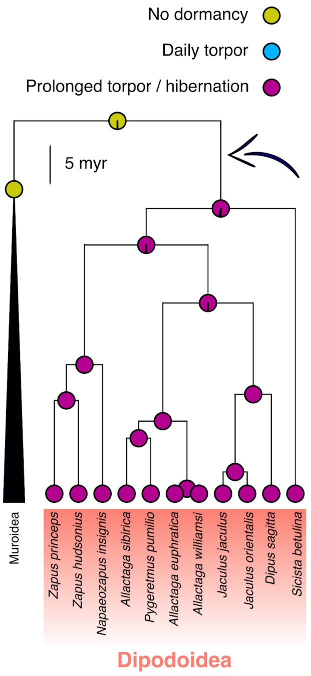


**Figure S8:** A direct shift between no dormancy and hibernation, denoted by an arrow, across the Myomorpha suborder. Muroidea and Dipodoidea are the two superfamilies that together comprise Myomorpha.

### S2. Ecophysiological data used in this study

#### S2.1. Sources of ecophysiological data for extant species

**Table S1:** The 21 ecophysiological variables collected for each species, along with their units and sources. For each variable, we progressively added data to our dataset by going through each source in the order shown here.

| Variable | Type & units / levels | Sources | Notes |
|---|---|---|---|
| Body mass | Continuous (g) | **1.** Genoud *et al.* (2018)<br>**2.** Smaers *et al.* (2021)<br>**3.** Tobias *et al.* (2022)<br>**4.** Herberstein *et al.* (2022)<br>**5.** Burger *et al.* (2019)<br>**6.** McNab (2008)<br>**7.** McNab (2009)<br>**8.** Ruf & Geiser (2015)<br>**9.** Nowack *et al.* (2020)<br>**10.** Nowack *et al.* (2023)<br>**11.** Font *et al.* (2019) | - |
| Basal metabolic rate | Continuous (W) | **1.** Genoud *et al.* (2018)<br>**2.** Londoño *et al.* (2015)<br>**3.** Herberstein *et al.* (2022)<br>**4.** McNab (2008)<br>**5.** McNab (2009) | For source #1, measurements were converted from ml $O_2 \cdot h^{-1}$ to W (1 W = 3,600 / 20.1 ml $O_2 \cdot h^{-1}$). For sources #4 and #5, measurements were converted from kJ $\cdot$ $h^{-1}$ to W (1 W = 3.6 kJ $\cdot h^{-1}$). |
| Brain mass | Continuous (g) | **1.** Herberstein *et al.* (2022)<br>**2.** Smaers *et al.* (2021)<br>**3.** Burger *et al.* (2019)<br>**4.** Font *et al.* (2019)<br>**5.** Jiménez-Ortega *et al.* (2020) | - |
| Maximum longevity | Continuous (yr) | **1.** Tacutu *et al.* (2013) | We only included data points with an "acceptable" or "high" quality. |
| Migration | Categorical (no / yes) | **1.** Webber & McGuire (2022)<br>**2.** Gnanadesikan *et al.* (2017)<br>**3.** Hardesty-Moore *et al.* (2018)<br>**4.** Tobias *et al.* (2022)<br>**5.** McNab (2009) | We treated partial migrants as migratory. |
| Carnivory | Categorical (no / yes) | **1.** Clarke & O'Connor (2014)<br>**2.** Jones *et al.* (2009)<br>**3.** McNab (2008)<br>**4.** Tobias *et al.* (2022) | We treated a species as capable of carnivory if even a small part of its diet was non-herbivorous, following the "TroL2" classification of Clarke & O'Connor (2014). |
| Herbivory | Categorical (no / yes) | **1.** Clarke & O'Connor (2014)<br>**2.** Jones *et al.* (2009)<br>**3.** McNab (2008)<br>**4.** Tobias *et al.* (2022) | As above. |
| Fossoriality | Categorical (nonfossorial / semifossorial / fossorial) | **1.** Healy *et al.* (2014) | - |

| | | | |
|---|---|---|---|
| Diurnality | Categorical (no / yes) | **1.** Cox *et al.* (2021)<br>**2.** Healy *et al.* (2014)<br>**3.** Maor *et al.* (2017)<br>**4.** Stark *et al.* (2020) | For source #3, we excluded species if studies disagreed on their activity patterns. |
| Crepuscularity | Categorical (no / yes) | **1.** Cox *et al.* (2021)<br>**2.** Healy *et al.* (2014)<br>**3.** Maor *et al.* (2017)<br>**4.** Stark *et al.* (2020) | See above. |
| Nocturnality | Categorical (no / yes) | **1.** Cox *et al.* (2021)<br>**2.** Healy *et al.* (2014)<br>**3.** Maor *et al.* (2017)<br>**4.** Stark *et al.* (2020) | See above. |
| Cathemerality | Categorical (no / yes) | **1.** Cox *et al.* (2021)<br>**2.** Healy *et al.* (2014)<br>**3.** Maor *et al.* (2017)<br>**4.** Stark *et al.* (2020) | See above. |
| Aquatic affinity | Categorical (very low / low / moderate / high) | **1.** Healy *et al.* (2014)<br>**2.** Pap *et al.* (2020)<br>**3.** Tobias *et al.* (2022) | We defined aquatic affinity as follows: **high:** the animal lives nearly exclusively in water; **moderate:** the animal may swim, dive, or float on water in its natural environment; **low:** the animal may wade in water or choose habitats close to water sources; **very low:** none of the above. |
| Range size | Continuous ($km^2$) | **1.** Jones *et al.* (2009)<br>**2.** Tobias *et al.* (2022) | - |
| Absolute latitude at the range midpoint | Continuous (°) | **1.** Jones *et al.* (2009)<br>**2.** Ruf & Geiser (2015)<br>**3.** Webber & McGuire (2022)<br>**4.** Nowack *et al.* (2023)<br>**5.** Tobias *et al.* (2022) | - |
| Hemisphere at the range midpoint | Categorical (southern / northern) | **1.** Jones *et al.* (2009)<br>**2.** Ruf & Geiser (2015)<br>**3.** Webber & McGuire (2022)<br>**4.** Nowack *et al.* (2023)<br>**5.** Tobias *et al.* (2022) | - |
| Mean temperature at the range midpoint | Continuous (°C) | **1.** Karger *et al.* (2017) | The "`bio1`" variable of CHELSA v2.1. |
| Temperature seasonality at the range midpoint | Continuous (°C) | **1.** Karger *et al.* (2017) | The "`bio4`" variable of CHELSA v2.1, divided by 100. |
| Annual precipitation at the range midpoint | Continuous ($kg \cdot m^{-2} \cdot yr^{-1}$) | **1.** Karger *et al.* (2017) | The "`bio12`" variable of CHELSA v2.1. |
| Precipitation seasonality at the range midpoint | Continuous ($kg \cdot m^{-2}$) | **1.** Karger *et al.* (2017) | The "`bio15`" variable of CHELSA v2.1. |
| Net primary productivity at the range midpoint | Continuous ($g\ C \cdot m^{-2} \cdot yr^{-1}$) | **1.** Karger *et al.* (2017) | The "`npp`" variable of CHELSA v2.1. |

#### S2.2. Sources of ecophysiological data for internal tree nodes

##### S2.2.1. Body mass

We collected body mass estimates for a few deep nodes in our phylogeny from previously published ancestral reconstructions based on fossil data. Specifically, we set the body mass of the last common ancestor of a) Placentalia to 170 g (Bertrand *et al.* 2022), b) Palaeognathae to 15,700 g (Torres *et al.* 2021), c) Neognathae to 2,900 g (Torres *et al.* 2021), d) Galloanserae to 3,050 g (Torres *et al.* 2021), and e) Neoaves to 1,450 g (Torres *et al.* 2021). For the last common ancestor of Amniota, Brocklehurst *et al*. (2022) did not provide an exact estimate but reconstructed it as being below 1 kg. Thus, we set its body mass to 2, 500, or 1,000 g as described in the main text.

##### S2.2.2. Brain mass

Similarly to body mass, we collected endocranial volume estimates from Bertrand *et al.* (2022) and Torres *et al.* (2021) for the last common ancestor of a) Placentalia (1.17 ml), b) Aves (7 ml), c) Palaeognathae (9.6 ml), d) Neognathae (5.6 ml), e) Galloanserae (5.9 ml), and f) Neoaves (5.4 ml). To convert endocranial volume estimates to brain mass estimates, we multiplied the former by a density value of 1.036 $g \cdot ml^{-1}$ (e.g., see Iwaniuk & Nelson 2002; Taylor *et al.* 1995).

##### S2.2.3. Basal metabolic rate

Wiemann *et al.* (2022) showed that it is possible to infer the metabolic rate of fossils from advanced lipoxidation end-product signals which are preserved during fossilization. They then reconstructed the evolution of mass-specific metabolic rate across a phylogeny of amniotes. From their ancestral reconstruction, we selected three nodes that had many descending lineages and, therefore, their estimates are likely to be quite robust: the last common ancestors of a) Amniota (2.08 $ml\ O_2 \cdot h^{-1} \cdot g^{-1}$), b) Placentalia (7.68 $ml\ O_2 \cdot h^{-1} \cdot g^{-1}$), and c) Neognathae (5.35 $ml\ O_2 \cdot h^{-1} \cdot g^{-1}$). Next, we multiplied these estimates by the reconstructed body masses of these three taxa (see above) to get mass-independent metabolic rate, and converted them to W units. Nevertheless, because of the experimental approach of Wiemann *et al.* (2022), the resulting metabolic rate estimates should be closer to field metabolic rate rather than basal metabolic rate. According to Clarke (2017), the ratio of field metabolic rate to basal metabolic rate is 2.37 for reptiles ($N$ = 33), 3.53 for mammals ($N$ = 18), and 3.44 for birds ($N$ = 13). Thus, we divided the metabolic rate estimates of Placentalia and Neognathae by 3.53, and 3.44, respectively, resulting in basal metabolic rate estimates that were close to the predicted values for extant endotherms of similar body size. For the last common ancestor of Amniota, we divided its metabolic rate value by 2.8, a value intermediate to those of reptiles (ectotherms) and mammals and birds (endotherms), but closer to that of the ectothermic group.

##### S2.2.4. Carnivory and herbivory

According to Clack (2012), Grossnickle *et al*. (2019), and O'Leary *et al*. (2013), there is strong evidence that the last common ancestors of a) Amniota, b) Mammalia, c) Theria, and d) Placentalia had non-herbivorous diets. Therefore, we set the corresponding nodes in our phylogeny as capable of carnivory and incapable of herbivory.

##### S2.2.5. Aquatic affinity

The last common ancestor of Amniota lived in coal swamps (Clack 2012) and, thus, we set its aquatic affinity to either low or moderate (see main text).

### S3. Additional MCMCglmm specification details

#### S3.1. Continuous imputation of missing values in response variables

Given that most of our response variables contained missing values for some species, we used the "missing at random" approach (Hadfield & Nakagawa 2010; de Villemereuil & Nakagawa 2014), implemented in `MCMCglmm`. In this approach, missing values in a response variable were continuously estimated at each step of the Markov chain, based on a combination of the phylogeny, known values for other species, and other co-varying response variables. This process yields unbiased estimates of missing values, as long as missingness is not systematically driven by a variable that is not present in the model.

#### S3.2. Variance-covariance matrices of response variables

Because we applied a phylogenetic random effect on the intercept of each response variable, the models estimated two separate variance-covariance matrices, one for variance captured by the phylogeny and one for the residual variance. We then summed the two matrices to obtain the total ("phenotypic") variance-covariance matrix. From this, we calculated pairwise correlation estimates by dividing the covariance of each pair of response variables by the square root of the product of their variances.

#### S3.3. Priors

We specified relatively uninformative priors, i.e., a parameter-expanded prior for the variance-covariance matrix of the phylogenetic random effect, an inverse gamma prior for the residual variance-covariance matrix, and the default normal prior for the intercept of each response variable. Because the model included threshold response variables, the total amount of variance was non-identifiable. Thus, following the `MCMCglmm` documentation, we fixed the residual variance to 22, i.e., the number of responses in the model.

#### S3.4. Number of generations and posterior diagnostics

To ensure that the posterior was sufficiently sampled and that convergence was reached, we executed 5 independent chains for each of the 6 models (30 chains in total; see Methods), for 1 million generations each. Each chain ran for an average of 7.5 days on a single AMD EPYC 7702 CPU core and required 67 GB of memory. Samples from the posterior were obtained every 25 generations after the first 10% (100,000 generations) which were discarded as burn-in. For each set of 5 chains, we calculated the effective sample size and the potential scale reduction factor. We verified that these were greater than 400 and smaller than 1.1, respectively, for all model parameters, indicating sufficient sampling and convergence (Brooks & Gelman 1998; Gelman & Rubin 1992). Lastly, we combined posterior samples across all 30 chains. We followed the same procedure for estimating correlations separately for mammals and for birds, with the only difference being that we executed chains for 2 million generations, sampling every 75 generations after a burn-in of 200,000 generations, which enabled chains to explore the parameter space sufficiently.

#### S3.5. Body mass-correction of metabolic rate, brain mass, and maximum longevity

We calculated the body mass-scaling exponents of these three traits as the covariance between body mass and the trait of interest (with both traits in log scale), divided by the variance of the natural logarithm of body mass, which were estimated by our `MCMCglmm` fits. We then divided the values of the three traits (in linear scale) by body mass raised to the median posterior value of the corresponding mass-scaling exponent.

### S4. Comparisons of the predictions of MCMCglmm fits with data

`MCMCglmm` fits generally predicted well the dormancy data (Supporting Fig. S9) and the 21 ecophysiological variables (Supporting Figs. S10 and S11). A notable exception was the dormancy capabilities of bears (Supporting Fig. S9B,D). `MCMCglmm` fits predicted daily torpor as the most likely state for all bears in our dataset (*Melursus ursinus*, *Ursus americanus*, *Ursus arctos*, and *Ursus maritimus*), whereas the observed state is lack of dormancy for *Melursus ursinus* and hibernation for the other species. Bears also had one of the few direct transitions between lack of dormancy and hibernation according to M*k* fits (Supporting Fig. S4). It is worth mentioning here that whether any bear species qualifies as a true hibernator remains under debate, given that they decrease their body temperature to only around 30°C (Evans *et al.* 2016; Hissa 1997; Tøien *et al.* 2011). Nevertheless, *Ursus americanus*, *arctos*, and *maritimus* have been observed to respond to environmental challenges by entering dens, where they minimise their activity and greatly reduce their metabolic rate ("metabolic denning"; Fowler *et al.* 2021). For this reason, we treated the aforementioned bear species as hibernators in our analyses.

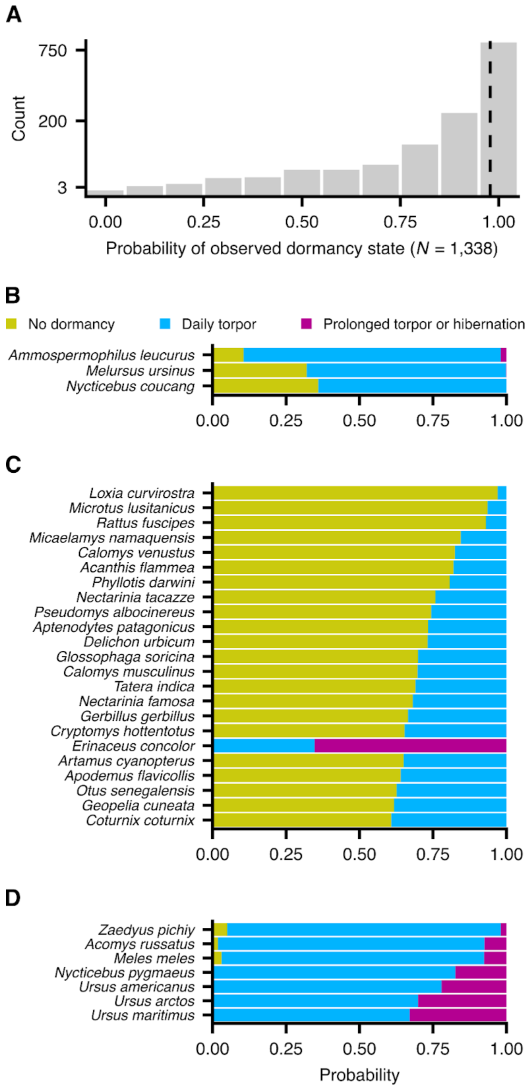


**Figure S9:** Comparison of the predictions of `MCMCglmm` fits for dormancy against our data. Panel A shows the posterior probabilities for the dormancy state listed in our dataset (observed state) per species. Posterior probabilities of 1 and 0 indicate complete agreement and disagreement, respectively. The dashed line stands for the median posterior probability across all species. Panels B-D show species for which the observed dormancy state (no dormancy, daily torpor, and hibernation, respectively) had a posterior probability below 0.4.

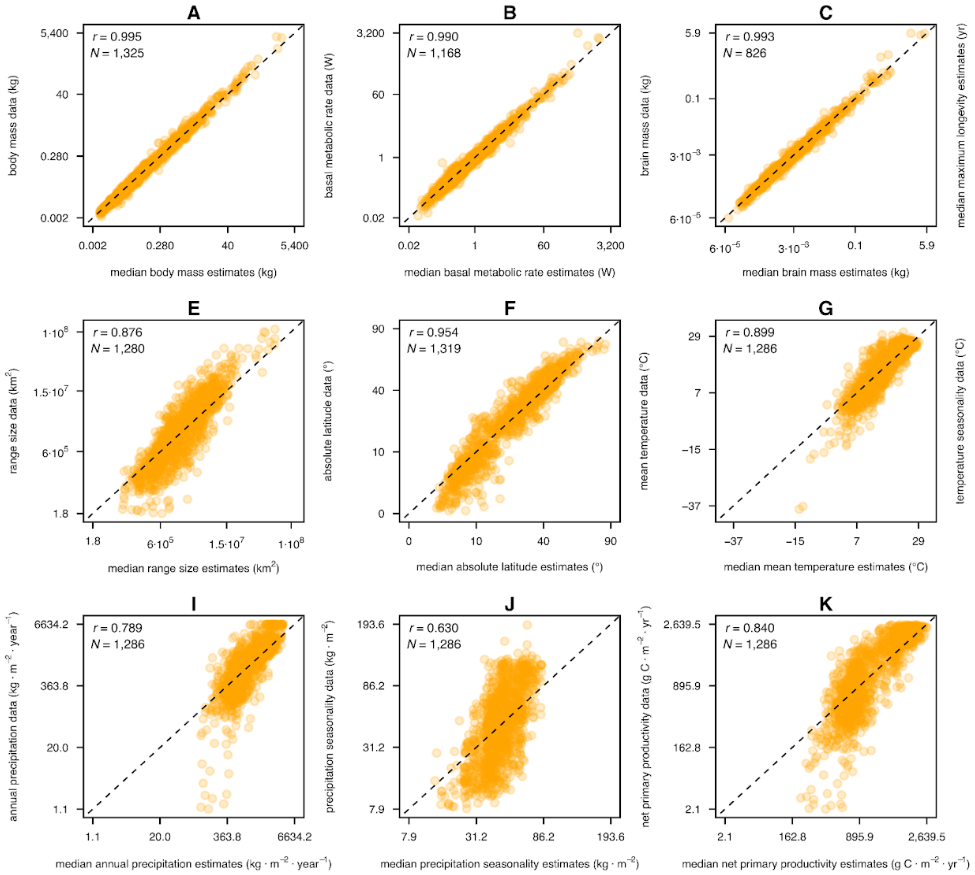


**Figure S10:** Scatterplots of the median predictions of `MCMCglmm` fits (horizontal axes) against the corresponding values in our dataset (vertical axes) for continuous ecophysiological variables of extant species. Dashed lines are the one-to-one lines. For most variables (especially those with strong phylogenetic signal), model predictions are well in line with the data, given the high correlation coefficient values reported for each pair of axes.

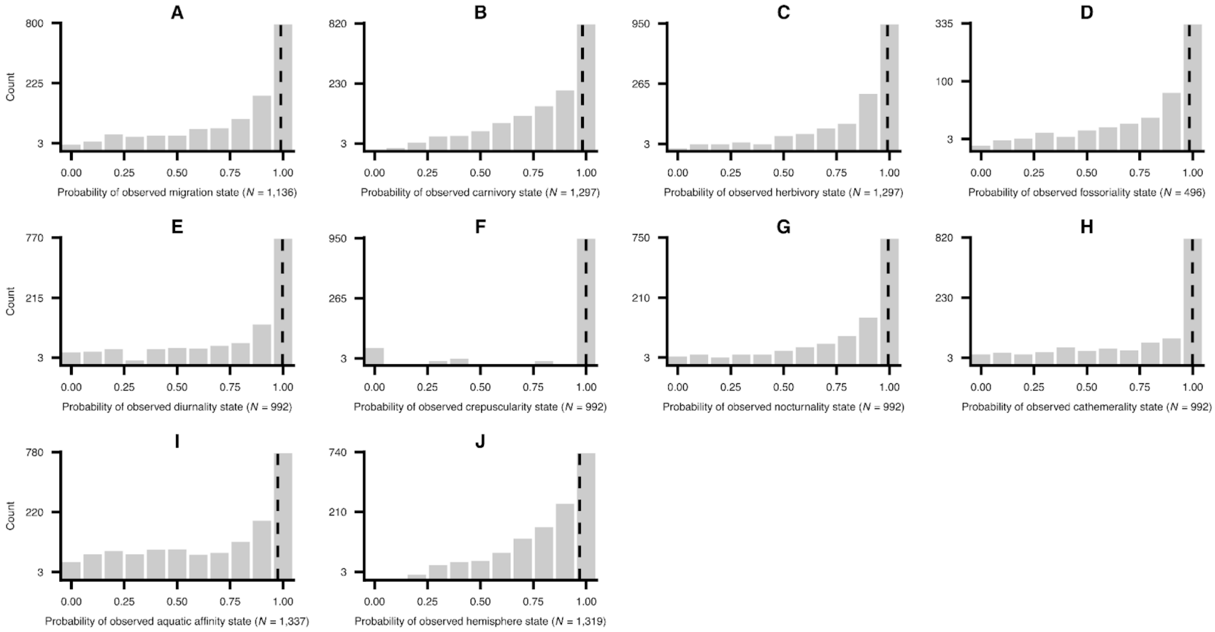


**Figure S11:** Comparison of the predictions of `MCMCglmm` fits for categorical variables in our dataset. Each panel shows the posterior probabilities for the state listed in our dataset (observed state) per species and variable. Posterior probabilities of 1 and 0 indicate complete agreement and disagreement, respectively. Dashed lines stand for the median posterior probability per variable.

#### S5. The relationship between migration and body mass in endotherms

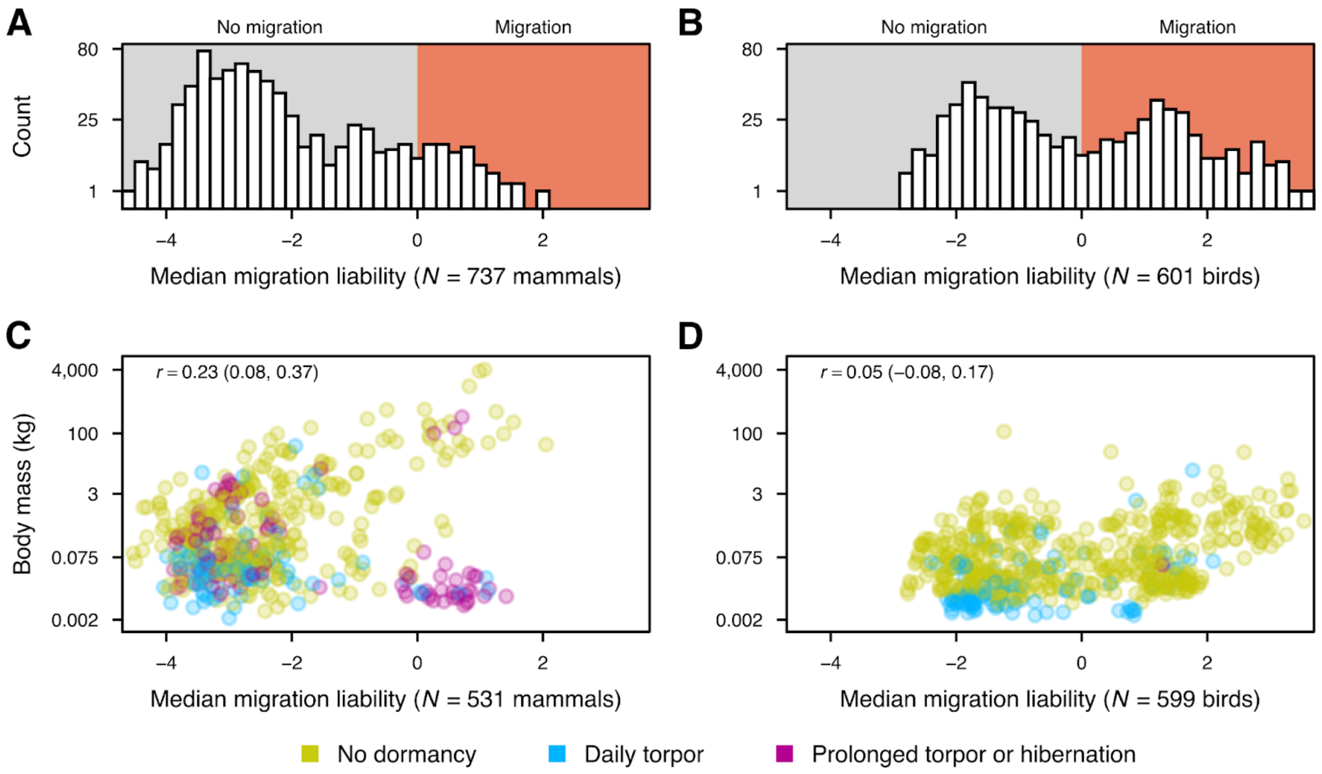


**Figure S12:** Migration and body mass in extant mammals (A, C), and birds (B, D). Panels A and B show the median liabilities for migration for all mammals and birds, respectively, in the present study. Panels C and D show body mass against median migration liability, only for species without missing values in these two variables. The median posterior correlation estimates between migration and the natural logarithm of body mass are explicitly reported, along with their 95% HPD intervals in parentheses. Across mammals (panel C), migration has a weak positive correlation with body mass, with bats being key outliers (cluster of data points with a migration liability near 0 or higher and a body mass below 3 kg). In contrast, across birds (panel D), migration is independent of body mass. Note that the values along the vertical axes of panels C and D do not increase linearly.

#### S6. Further analyses of the distribution of dormancy along the ecophysiological parameter space

##### S6.1. Projection of dormancy descriptors onto the ecophysiological parameter space

To understand if slight differences in dormancy characteristics are linked with distinct areas of the ecophysiological parameter space (see Figs. 4 and 5 in the main text), we used three dormancy descriptors for mammalian species introduced by Nowack *et al.* (2023): i) seasonality (whether dormancy occurs in only a single season), ii) predictability (whether conspecifics tend to enter dormancy in a similar manner), and iii) preparation for hibernation. Based on these, we found that different subtypes of dormancy tend to overlap substantially (rather than segregate) across the ecophysiological parameter space (Supplementary Figs. S13-S15).

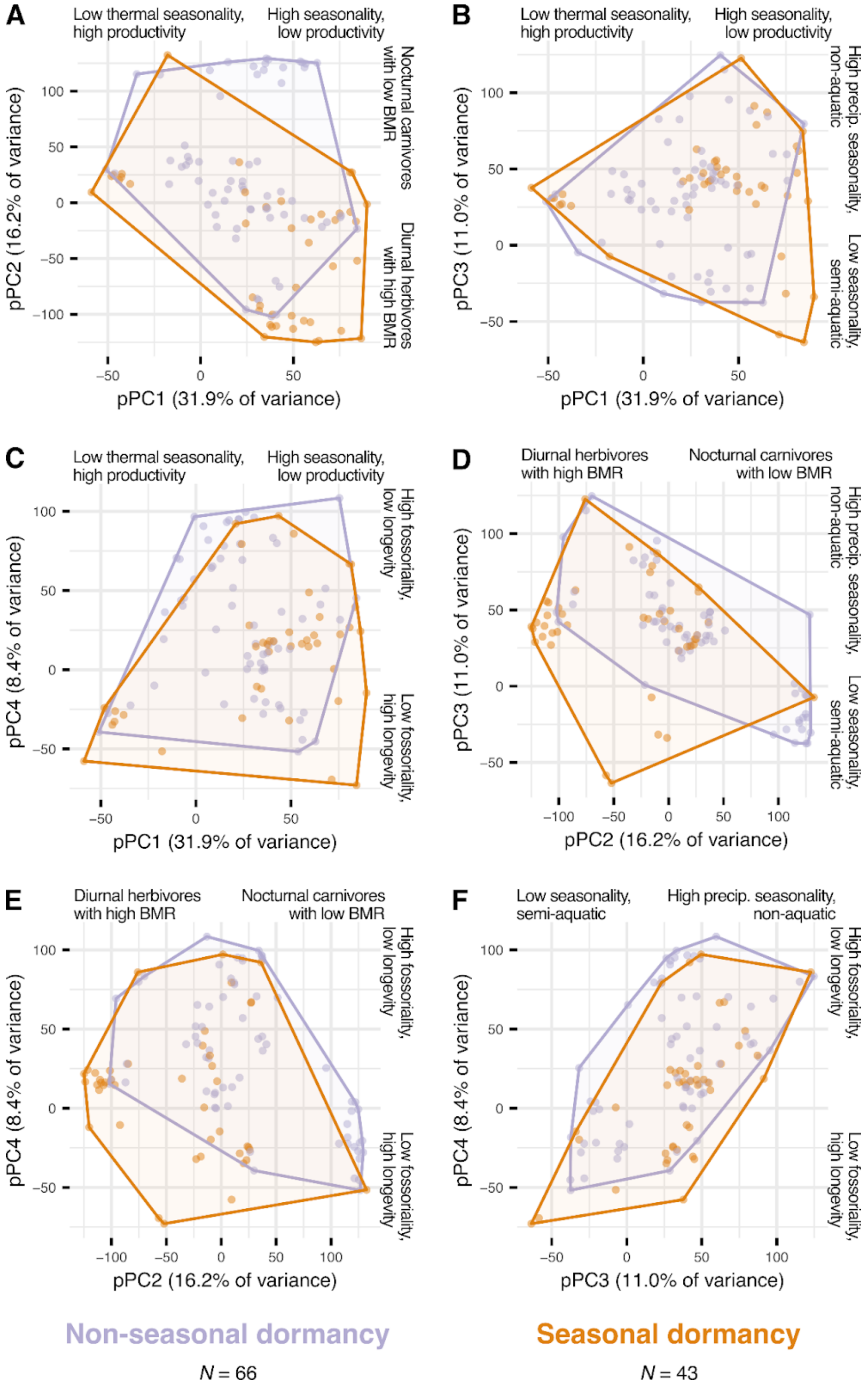


**Figure S13:** Distribution of dormancy seasonality along the ecophysiological parameter space. Data points stand for dormancy-capable mammals.

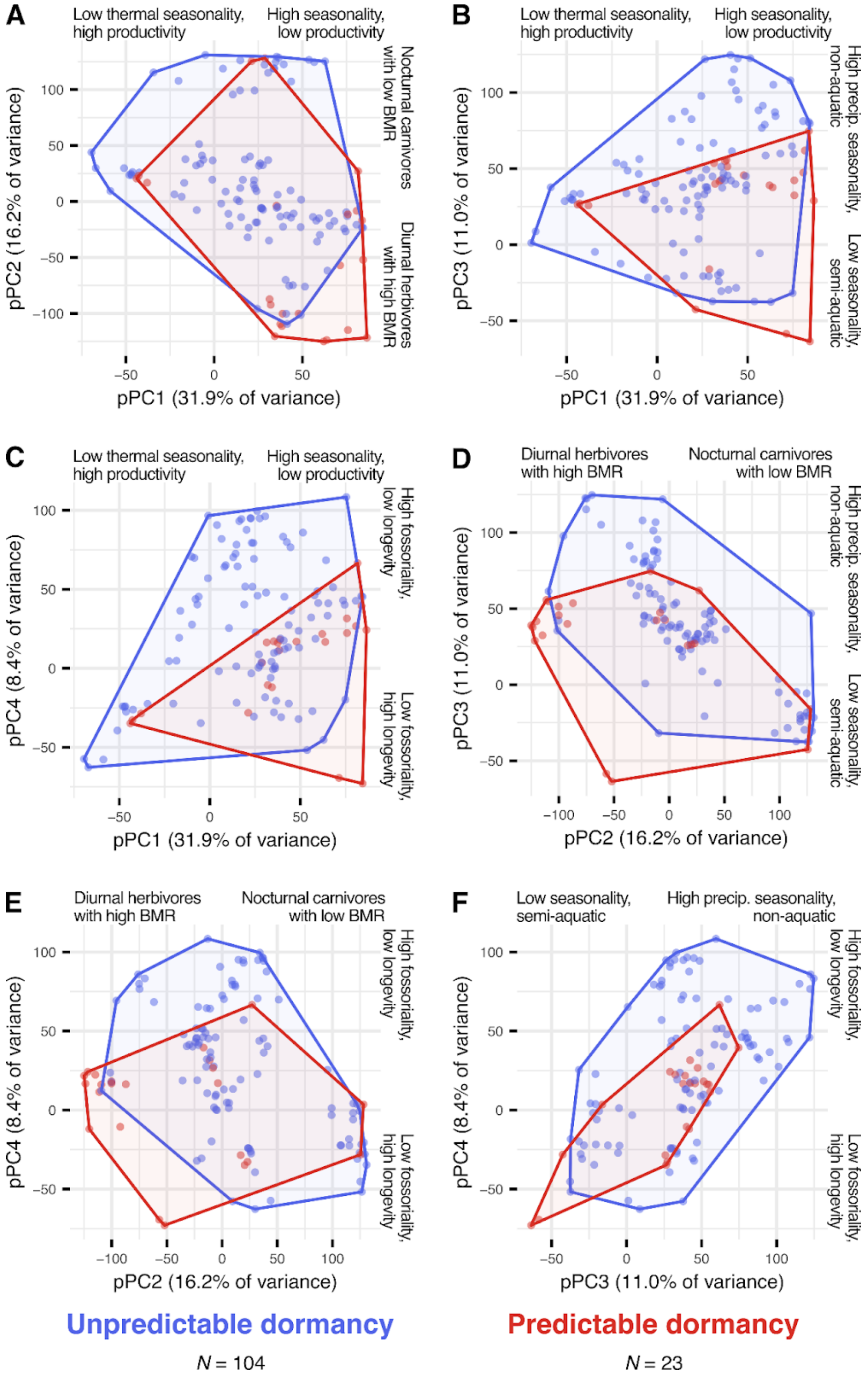


**Figure S14:** Distribution of dormancy predictability along the ecophysiological parameter space. Data points stand for dormancy-capable mammals.

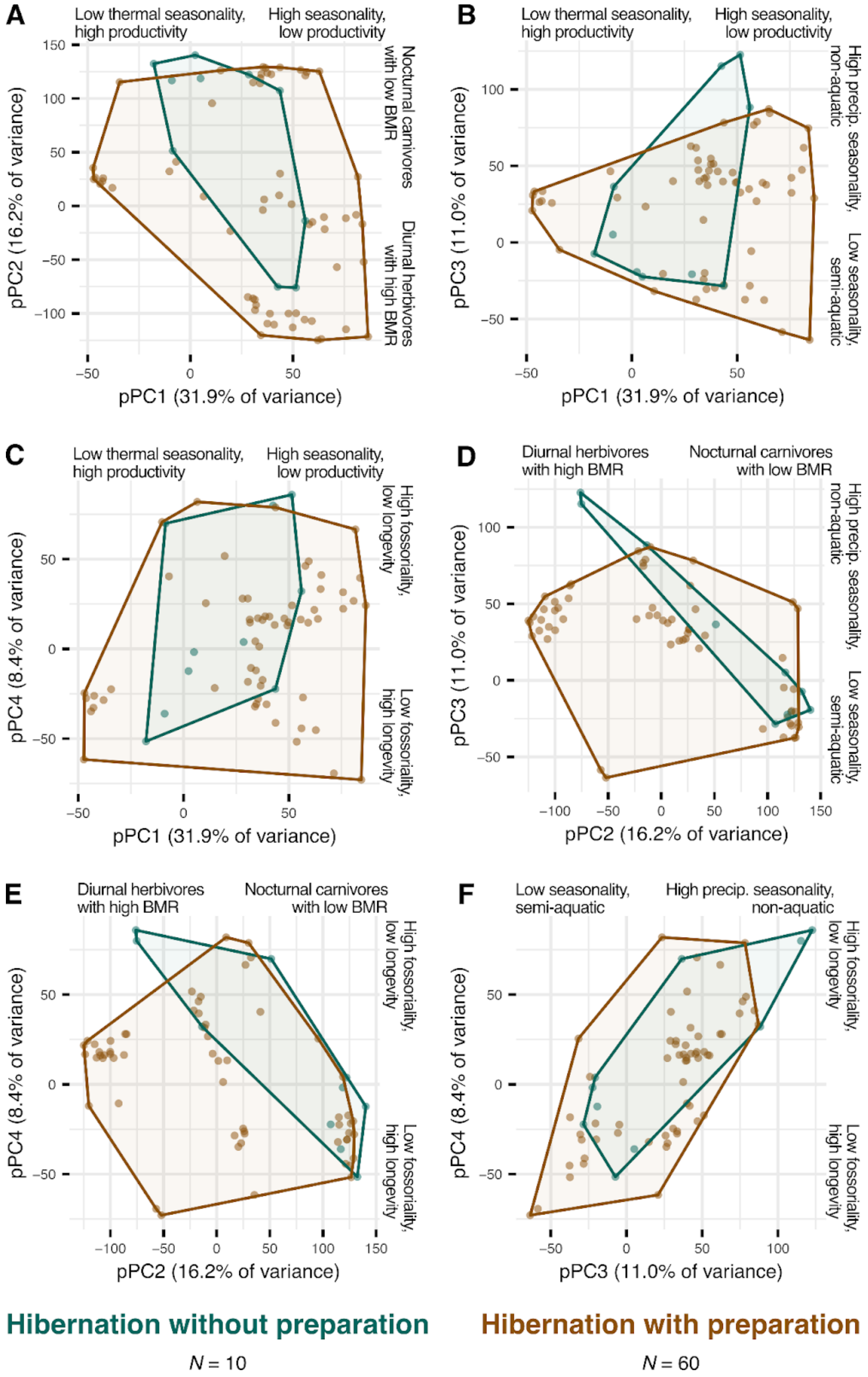


**Figure S15:** Distribution of hibernation preparation (or lack thereof) along the ecophysiological parameter space. Data points stand for mammalian hibernators.

#### S6.2. Phylogenetic PCA across only mammal species

We additionally applied a phylogenetic PCA to only dormancy-capable mammals in our dataset to investigate if the resulting patterns differ from those obtained when we analysed birds and mammals simultaneously (Fig. 4 in the main text). Performing the analysis separately for mammals led to qualitatively identical results (Supplementary Fig. S16).

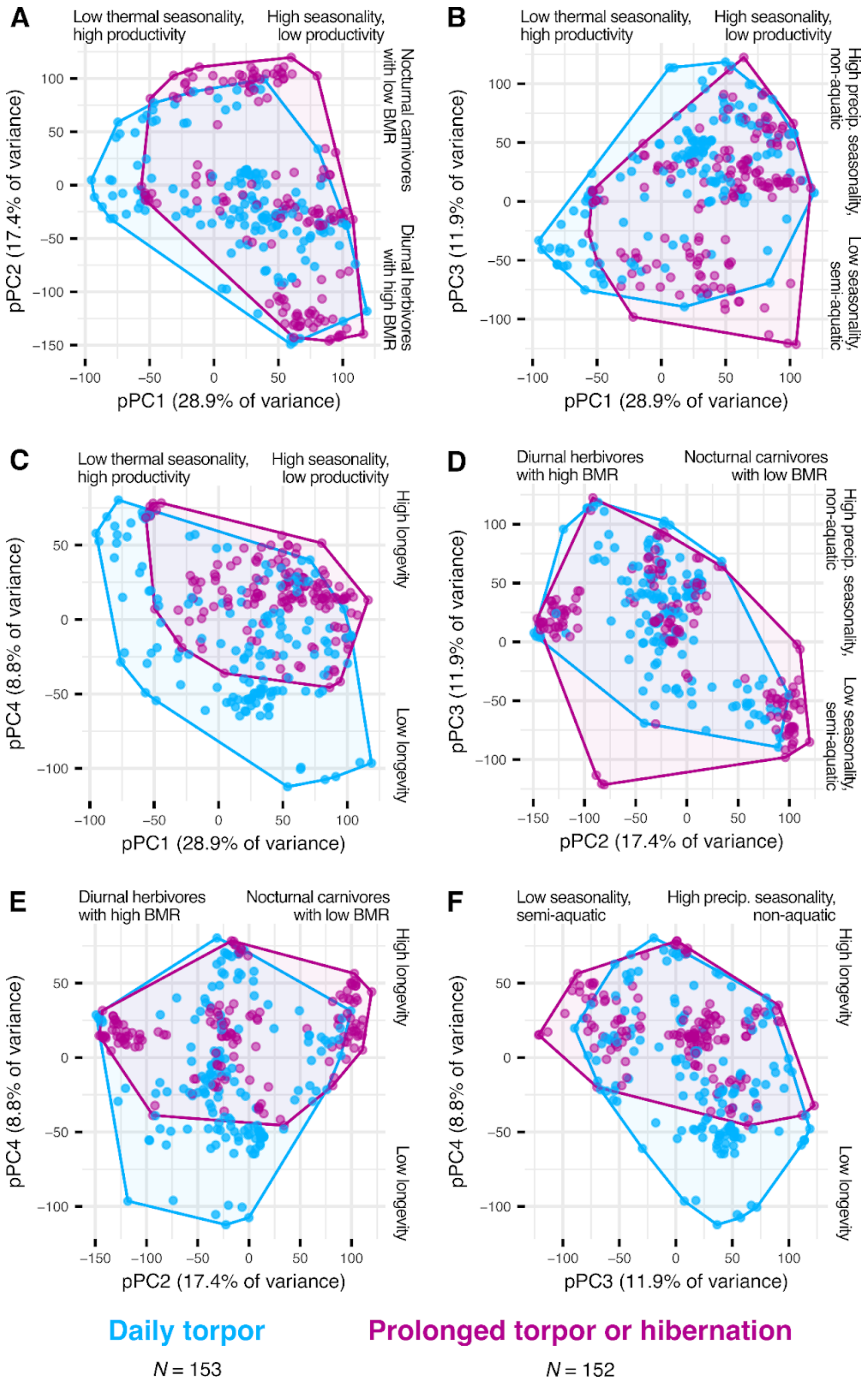


**Figure S16:** Distribution of daily torpor and hibernation along the ecophysiological parameter space, estimated by including only mammalian dormancy-capable species in the phylogenetic principal components analysis.
